# Distinct sub-second dopamine signaling in dorsolateral striatum measured by a genetically-encoded fluorescent sensor

**DOI:** 10.1101/2022.01.09.475513

**Authors:** Armando G. Salinas, Jeong O. Lee, Shana M. Augustin, Shiliang Zhang, Tommaso Patriarchi, Lin Tian, Marisela Morales, Yolanda Mateo, David M. Lovinger

**Affiliations:** Laboratory for Integrative Neuroscience, National Institute on Alcohol Abuse and Alcoholism, National Institutes of Health, Rockville, MD, USA; Department of Bioengineering, George Mason University, Fairfax, VA, USA; Confocal and Electron Microscopy Core, National Institute on Drug Abuse, Baltimore, MD, USA; Department of Biochemistry and Molecular Medicine, University of California at Davis, Davis, CA, USA; Institute of Pharmacology and Toxicology, University of Zurich, Zurich, Switzerland; Neuronal Networks Section, Integrative Neuroscience Research Branch, National Institute on Drug Abuse, Baltimore, MD, USA; Current affiliation, Department of Pharmacology, Toxicology & Neuroscience, Louisiana State University Health Sciences Center – Shreveport, Shreveport, LA, USA; Current affiliation, Department of Pharmacology, Northwestern University Feinberg School of Medicine, Chicago, Illinois, USA

## Abstract

Dopamine produces neuromodulation throughout the basal ganglia, cortex and other brain regions, and is implicated in movement control, neural mechanisms of reward and actions of misused substances. The efferent projections of dopaminergic neurons with somata in the substantia nigra pars compacta and ventral tegmental area strongly innervate different striatal subregions. While much is known about the function of these neurons, there is a relative deficiency of information about *in vivo* dopamine dynamics in the nigrostriatal projections, especially those to the dorsolateral striatum (DLS). In past studies, subsecond dopamine changes were measured predominantly with fast-scan cyclic voltammetry (FSCV) both in brain slices and *in vivo*. However, traditional FSCV has limitations in discriminating among catecholamines, and cannot be used for simultaneous measurement of both slow and fast/phasic dopamine changes. In addition, FSCV has been most useful for measuring dopamine in the ventral striatum *in vivo* with less utility for measurement in dorsolateral striatum. The development of genetically encoded dopamine sensors has provided a new approach to measuring slow and fast dopamine dynamics both in brain slices and *in vivo*, raising the hope of more facile measurement of *in vivo* dopamine measurements, including in areas where measurement was previously difficult with FSCV. To this end, we first evaluated dLight photometry in brain slices with simultaneous FSCV. We found that both techniques yielded comparable findings. However, differences were noted in responses to dopamine transporter inhibitors, including cocaine. We then used *in vivo* fiber photometry with dLight to examine responses to cocaine in DLS and compared responses during Pavlovian conditioning in DLS to two other striatal subregions. These experiments show that dopamine increases are readily detectable in DLS and provide new information about dopamine transient kinetics and slowly developing signaling during conditioning. Overall, our findings indicate that dLight photometry is well suited to measuring dopamine dynamics in a striatal region of great interest where such measurements were difficult previously.

## Introduction

Dopamine is a catecholamine neurotransmitter found throughout the mammalian nigrostriatal and cortical-mesolimbic circuit with critical roles in many psychiatric and neurological disorders including Parkinson’s Disease, psychosis, schizophrenia, and addiction (e.g. Nestler and Carlezon, 2006; Calabresi et al., 2009; Grace, 2016; Berke, 2018; Ott and Nieder, 2019). Midbrain dopaminergic neurons provide extensively branching axons to the striatum. The dopamine released from these afferents produces neuromodulation with key roles in neuronal function, synaptic plasticity and behavior. The rodent striatum has several subregions that are part of associative (dorsomedial striatum, DMS), limbic (nucleus accumbens/ventral striatum, NAc) and sensorimotor (dorsolateral striatum, DLS) circuits that have different roles in learning, movement control and reward (Yin and Knowlton, 2004). Thus, understanding the dynamics of dopamine release in different striatal subregions is important for a full understanding of the function of these striatal regions in the context of different brain circuits.

Dopamine release has been measured in isolated cells, brain slices and *in vivo* with techniques including microdialysis sampling coupled with electrochemical detection and variants of fast-scan cyclic voltammetry (FSCV). Microdialysis allows for precise chemical identification and concurrent measurements of multiple neurotransmitters from each sample (Chefer et al., 2009). Variants of microdialysis can also be used to estimate the absolute dopamine concentration in a brain region. However, this technique has a relatively slow sampling time, typically 5-20 minutes per sample that does not allow for precise correlation of dopamine levels with discrete behavioral events. Fast-scan cyclic voltammetry is an electrochemical method that can be used to measure phasic release of catecholamines at sampling rates of 10Hz or higher (Venton and Cao, 2020). Thus, FSCV allows for real-time measurement of dopamine release in response to stimuli in brain slices, as well as *in vivo* detection of sub-second changes in neurotransmitter release in relation to behavioral events. Indeed, real-time dopamine measurements with FSCV revolutionized the study of fast dopamine changes and motivated behaviors for well over a decade. However, FSCV has limitations. Conventional FSCV discerns fast signals from a recent baseline or pre-stimulus period, and thus this technique cannot measure absolute dopamine concentrations and is not well suited to measurement of slow changes in dopamine levels. In addition, FSCV cannot distinguish between different catecholamine neurotransmitters. In practice, FSCV measurements of *in vivo* dopamine have been made mainly in the rat Nucleus accumbens, although a few studies have examined the rat DMS and DLS (Brown et al., 2011; Howe et al., 2013; Klanker et al., 2017; Willuhn et al., 2012). Unfortunately, *in vivo* measurements in mouse dorsal striatum, and the DLS in particular, are lacking. This is presumably due to difficulties in detecting small changes accompanied by contaminating signals.

Genetically encoded fluorescent biosensors for dopamine were recently developed (Patriarchi et al., 2018; Sun et al., 2018). These membrane-targeted, G-protein-coupled receptor (GPCR)-based sensors employ modified dopamine receptors inactive for intracellular signaling that have had a circular permuted Green Fluorescence Protein (cpGFP) molecule in place of the third intracellular loop. The sensors work by coupling the dopamine induced conformational changes to changes in cpGFP emission intensity. Although fluorescent dopamine sensors cannot measure absolute concentrations of dopamine, they can be used to simultaneously detect slow and fast (or phasic) changes in dopamine relative to baseline levels (Mohebi et al., 2019 ; Jørgensen et al. 2023; Van Zessen et al. 2021). These sensors also have the sensitivity to measure dopamine in brain regions where such measurements were previously difficult (e.g. cerebral cortex Patriarchi et al., 2018). Using genetically encoded fluorescent dopamine sensors allows for the real-time measurement of dopamine dynamics in relation to distinct behaviors in areas such as DLS.

In the current study we first evaluated the ability of the genetically encoded dopamine sensor dLight to detect stimulation-induced dopamine in dorsal striatum using brain slice photometry with simultaneous FSCV. We found many similarities in indices of regulation of dopamine release using both methods. We also noted differences in cocaine-induced effects on dorsal striatal dopamine release measured with dLight photometry and FSCV that challenge the tacitly accepted, but controversial, notion that cocaine enhances dopamine release *in vitro*. Next, we examined striatal dopamine dynamics using *in vivo* fiber photometry with dLight to assess slow and fast/phasic dopamine changes in DLS in comparison to other striatal subregions. In Pavlovian conditioning driven by a natural food reinforcer, we also found distinct characteristics of phasic dopamine dynamics in DLS compared to other striatal subregions. These findings provide new information about dopamine dynamics in dorsal striatum, and should promote further physiological and behavioral studies in dopamine dynamics throughout the striatum.

## Results

### Simultaneous dLight and FSCV dopamine measurement in DLS slices

We first wanted to examine the characteristics of dLight under conditions where dopamine is readily detected even in DLS. Thus, we performed simultaneous recordings using a PMT based photometer and traditional FSCV in acutely prepared brain slices (**Figure 1a**). Electrical stimulation evoked fluorescence transients were readily observed in DLS and were blocked by D1 dopamine receptor antagonism (**Figure 1b** **and Supplementary Figure 1**). Simultaneous dLight photometry and FSCV recordings in response to different stimulus intensities revealed similar input—output curves, with both methods showing significant responses to 100µA stimuli (**Figure 1c**, one sample t-test at 100µA, photometry t=5.128 p=0.0022, FSCV t=2.512, p=0.0458, df=6).

**Figure1.**
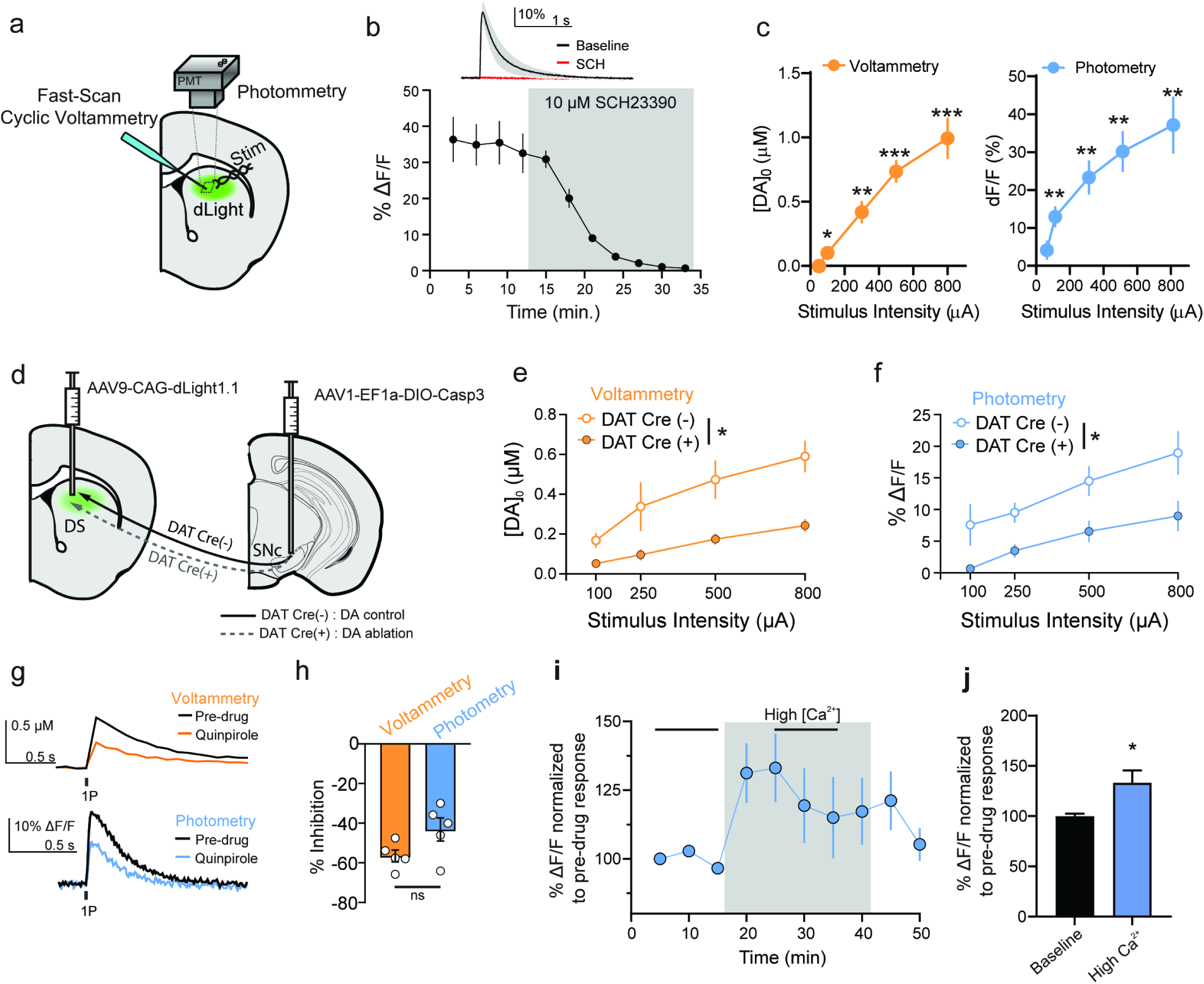
Simultaneouscomparison of dLight and FSCV dopamine responses in dorsal striatal slices. **(a)** Schematic diagram of simultaneous dLight photometric and FSCV recordings. **(b)** dLight signal is completely blocked by application of the D1 dopamine receptor antagonist, SCH23390 **(c)** Input-output curves of dLight photometric and FSCV DA measurements (*p<0.05, **p<0.01; ***p<0.005, one sample t-test). **(d)** Schematic diagram illustrating the viral strategy used to ablate substantia nigra DA neurons to confirm that changes in striatal dLight fluorescence are due to dopamine. **(e & f)** Genetic ablation of nigral DA neurons results in markedly reduced dopamine release measured with dLight and FSCV. Representative traces **(g)** and summarized data **(h)** showing that application of the D2 dopamine receptor agonist, quinpirole, inhibits dopamine release equally with both methods. **(i & j)** Increasing extracellular calcium increased dLight fluorescent responses. * p<0.05, ** p<0.01. ***p<0.001 Error bars represent the SEM.

### dLight fluorescence signals and FSCV originate from dopamine neurons and are dynamic

To ensure that the electrically evoked dLight fluorescence transients were attributable to dopamine release from midbrain neurons, we used a viral strategy to genetically ablate substantia nigra dopamine neurons. We infused a Cre-dependent Caspase3-encoding virus into the substantia nigra and dLight virus into the dorsal striatum of DAT Cre mice (**Figure 1d**). At least five weeks later, brain slices were prepared and simultaneous dLight and FSCV recordings were conducted. We found that genetic ablation of nigral DA neurons resulted in significantly reduced dopamine release in DAT Cre+ relative to DAT Cre mice, as measured with dLight (*, p<0.05, F(1,14)=5.257) and FSCV (*, p<0.05, F(1,7)=7.63) across several stimulation intensities **Figure 1e** **& f**). Also, the electrically evoked fluorescence signal was not detected in eGFP control experiments (**Supplementary Figure 2**)

We next assessed whether dLight and FSCV signals would show similar dynamics during manipulations that decrease or increase dopamine release. Thus, we applied the D2 dopamine receptor agonist, quinpirole, to slices to inhibit dopamine release (via activation of presynaptic D2 dopamine autoreceptors). We found that 30 nM quinpirole inhibited dopamine release equally with both methods (**Figure 1h**; t=2.032; p=0.076, df=8). Having shown that dLight signals could be decreased, we assessed whether they could be increased by increasing extracellular calcium levels, which would enhance dopamine release. Indeed, increasing extracellular calcium increased dLight fluorescent responses (**Figure 1i** **& j**; *, p<0.05, t=2.611, df=12) demonstrating that dLight signals are truly dynamic. Further, these results demonstrate comparable performance between dLight photometry and FSCV measurements in terms of sensitivity to presynaptic inhibition and excitability induced changed in dopamine release.

### Differential regulation of dLight photometry and FSCV dopamine signals by DAT inhibitors

There have been many reports of dopamine transporter (DAT) inhibitor-mediated increases in DA release published using FSCV. However, in Patriarchi et al. (2018), it appeared that cocaine did not increase the dLight dopamine transient peak amplitude while it increased transient duration. This intriguing observation directly conflicted with a large body of literature and prompted us to directly compare cocaine effects using dLight photometry and FSCV dopamine measurements. We began by comparing the effect of increasing cocaine concentrations on simultaneously collected dLight photometry and FSCV measurements (**Figure 2a**). We found that cocaine increased the peak amplitude of transients measured with FSCV, but not those measured with dLight photometry (**Figure 2b,c**). Cocaine increased the duration of both dLight photometry and FSCV-measured transients, suggesting that cocaine was in fact inhibiting DAT. Notably, cocaine concentrations greater than ∼10µM led to a decrease in peak transient amplitude measured with both techniques. This is consistent with previous findings using FSCV and is due to off target cocaine inhibition of nAChRs, which contribute significantly to striatal dopamine release. This off target cocaine inhibition of nAChRs was first shown electrophysiologically (Francis et al., 2000; Ma et al., 2020) and then with striatal FSCV (Acevedo-Rodriguez et al., 2014; Chen et al., 2019).

**Figure 2.**
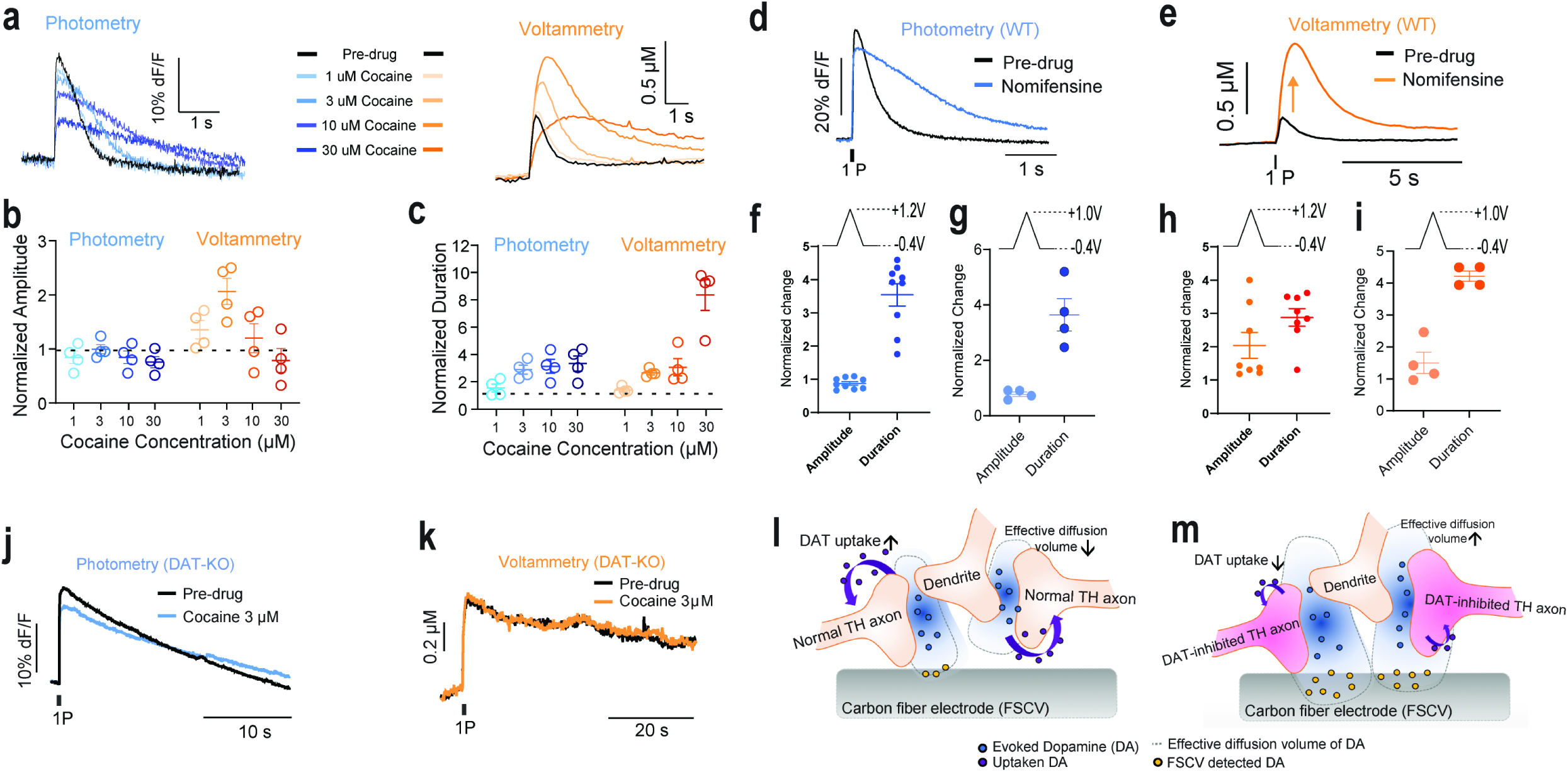
DAT blockers do not increase DA release. **(a)** Representative traces of simultaneously collected dLight (blue shades) and FSCV (orange shades) DA transients before and after application of cocaine. **(b)** Summary data showing that application of cocaine to dorsal striatal slices dose dependently increases peak DA transient peak amplitude measured with FSCV but not dLight photometry methods. **(c)** Summary data showing that DA transient duration is increased with both methods following cocaine application. **(d,e)** Similarly, the specific DAT inhibitor, nomifensine (1μM) increases DA transient peak amplitude with FSCV but not dLight photometry measurements. **(f)** Nomifensine experiment: Amplitude and duration of transients measured with dLight photometry normalized to pre-drug conditions with the unmodified triangle waveform (-0.4V to 1.2V) **(g)** The same normalized dlight photometry readout with nomifensine and modified triangle waveform (-0.4V to 1.0V). **(h)** The normalized FSCV amplitude and duration changes with nomifensine and unmodified triangle waveform (-0.4V to 1.2). **(i)** The normalized FSCV amplitude and duration changes with nomifensine and modified triangle waveform (-0.4V to 1.0V) **(j,k)** Cocaine does not increase DA transient peak height in DAT KO mice in photometry (j) and FSCV measurement (k). **(l,m)** Schematic diagram of the model for DAT inhibitor-induced increases in dopamine transient peak height measured using FSCV without DAT inhibitor(l) and with DAT inhbitor(m). Error bars represent the SEM.

We next determined whether a more specific DAT inhibitor, nomifensine (1 µM), would have effects similar to cocaine. Like cocaine, nomifensine increased the peak transient amplitude measured with FSCV (**Figure 2e,h**), but not with dLight photometry. Application of nomifensine increased the transient decay time measured with both techniques (**Figure 2d,f,h**).

### Altering FSCV adsorption altered DAT inhibitor effects on dopamine release

To further probe the factors underlying the DAT inhibitor-induced increase in peak transient amplitude measured with FSCV, we altered the triangle voltage wave form to applied to the carbon fiber electrode (CFE) to peak at +1.0V instead of +1.2V. (**Figure 2g,i**). This waveform change will decrease the dopamine aborption profile and effectively decrease the sensitivity of the FSCV CFE (Heien et al., 2003; Venton and Cao, 2020). As expected, nomifensine (1 µM) effects measured with dLight photometry were not affected by the waveform modification (**Figure 2f,g**). Note that the normalized amplitude of the fluorescence intensity was around 1, indicating that the dLight photometry measurement reports no increased dopamine release in response to the DAT inhibitor.

When we assessed evoked dopamine release before and after application of nomifensine (1 µM) using the modified triangle waveform for FSCV measurements, we found that DAT inhibition increased dopamine transient duration (**Figure 2h**) consistent with the dLight photometry measurement. However, no change in peak transient amplitude was detected during nomifensine application when the altered triangle waveform (peak at +1.0V) was used for FSCV measurements (**Figure 2i**). Thus, the apparent increase in dopamine release measured with FSCV cannot be detected if CFE absorption is decreased.

### DAT inhibitors do not affect dopamine release in DAT KO mice

FSCV has been used extensively to study the effects of cocaine and other uptake blockers on the dynamics of dopamine clearance and evoked release. Indeed, to explain how cocaine might lead to increased dopamine release (as measured with FSCV), it was posited that cocaine acted via a DAT-independent mechanism (Venton et al., 2006). To evaluate this hypothesis, we again performed simultaneous dLight photometry and FSCV recordings during which cocaine was applied to dorsal striatum brain slices from DAT KO mouse expressing dLight (**Figure 2j**). In DAT KO mice, the duration of stimulation-induced dopamine transients is prolonged compared to WT mice with a decay in the 10s of seconds, compared to 1-2s in WT mice (Jones et al., 1998; Budygin et al., 2002; Mateo et al., 2004). Accordingly, in our experiments, both dLight photometry and FSCV evoked dopamine transients were prolonged relative to WT mice (**Figure 2j,k**). Cocaine did not affect either dopamine transient amplitude or decay time measured with FSCV or dLight photometry. If the putative cocaine-induced increases in dopamine release assessed with FSCV are due to DAT-independent mechanisms, then the increase in FSCV dopamine transient peak height should be present in DAT KO mice. Since this was not the case, we considered other potential mechanisms for the putative DAT inhibitor-induced increases in dopamine release observed with FSCV.

### DAT inhibitors do not increase dopamine release: a new interpretation of results to reconcile differences with different methods

To reconcile these results, we posited a new interpretation for the effects of DAT inhibitors on dopamine transients measured with FSCV (**Figure 2l**). The detection of dopamine with FSCV relies on the electrochemical oxidation of dopamine at the surface of the CFE. Upon electrical stimulation of the slice, dopamine is released and diffuses across a given volume. The extent of this diffusion is determined by several factors but most notably DAT (Venton et al., 2003; Cragg and Rice, 2004). Thus, in a given space, the sampling volume obtained with FSCV is limited to the distance that dopamine can diffuse to the CFE surface. Under conditions of impaired DAT function, however, the diffusion distance of dopamine increases (**Figure 2m**). This increased release point diffusion of dopamine also increases the effective sampling volume of the CFE, resulting in a larger signal, but not increased release per se. When the dopamine absorption/sensitivity of the CFE is reduced (e.g. with the altered triangle waveform, Figure 2g,i) the effect of the increased sampling volume is reduced and thus the enhanced overflow does not alter peak transient amplitude.

### dLight sensor traffics to synaptic and extrasynaptic sites

We next sought to determine if dLight could traffic to synaptic sites. This would mark a major advance in the measurement of dopamine as both microdialysis and FSCV can only measure non-synaptic dopamine overflow. We performed immunohistochemistry for tyrosine hydroxylase (TH), the rate limiting step in catecholamine synthesis and GFP to label dLight in mouse dorsal striatum. We found putative dopamine axons (i.e. TH+ axons) juxtaposed to dLight/GFP (**Figure 3a**), suggesting the possibility of dLight expression in close proximity to dopamine release sites. We followed up these experiments with electron microscopy (EM) experiments to directly assess whether dLight would traffic to synapses and thus, at least partially, report synaptic dopamine release (**Supplementary Table S1**). We found that dLight trafficked exclusively to plasma membrane (**Figure 3b**) or membrane-associated regions within the cell (**Figure 3c**). We also found that dLight could indeed traffic to synaptic sites, including putative dopamine synapses (**Figure 3d**), and was also present at nearby extrasynaptic sites ranging from a few μm to 10s of μm from synapses.

**Figure 3.**
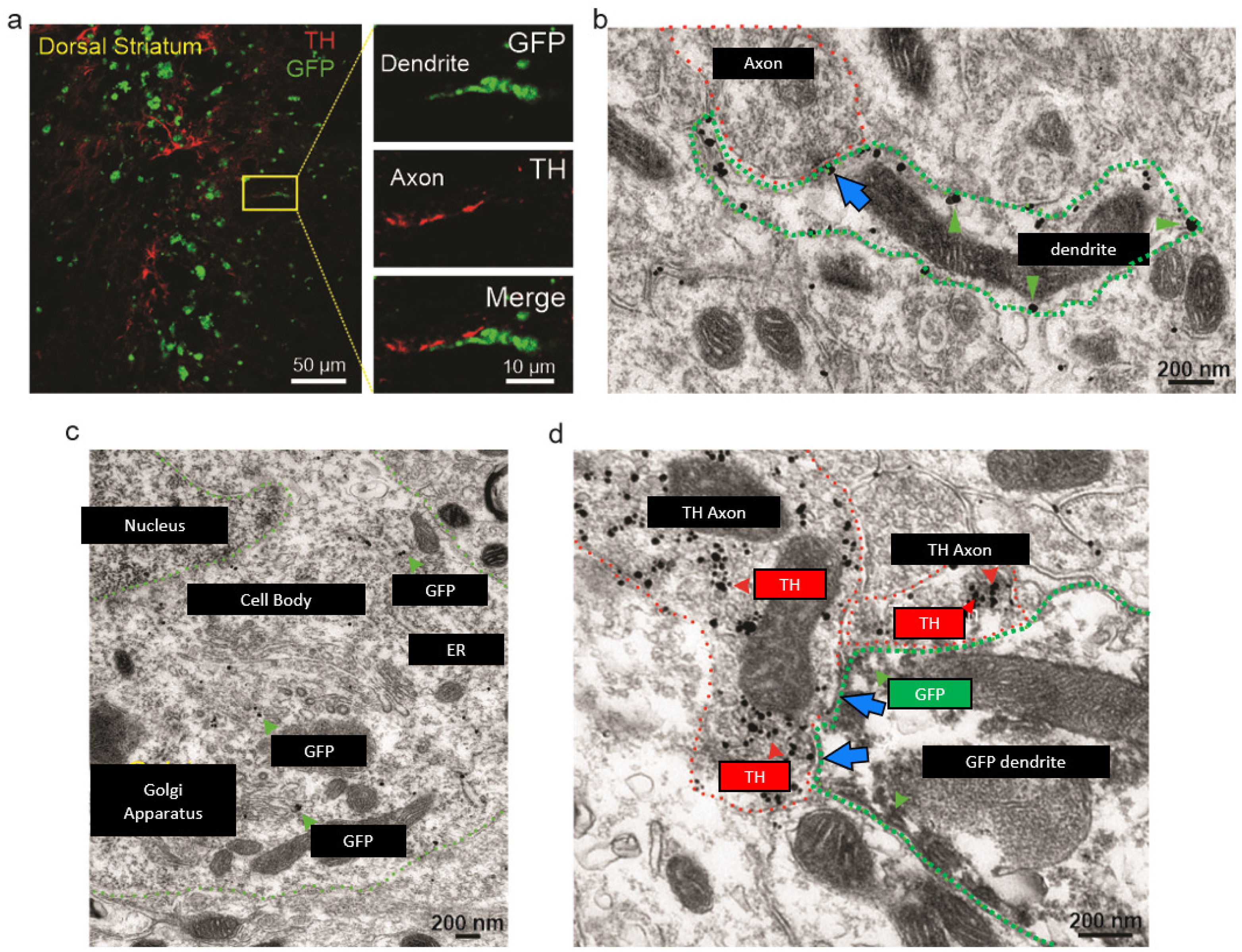
Dorsal striatum expression of the dLight sensor on plasma membrane of dendrites synapsing with TH-axons. Immunofluorescence detection of dLight sensor with an antibody targeted to the GFP moiety contained in the sensor (green) and tyrosine hydroxylase (TH, red) in the dorsal striatum from a mouse injected with a viral vector encoding dLight sensor. **(a)** Apposition of a dLight-positive dendrite and TH-positive axon is seen at high magnification (right panels). **(b)** Electron micrographs of immunogold detection of dLight sensor (gold particles, green arrowheads) on the plasma membrane of a dendrite (green outline in b). Note the apposition of dendritic dLigh and a presynaptic axon (blue arrow in b). **(c)** dLight expression on the plasma membrane of a soma (green outline). Note also the dLight sensor (gold particles) in association with Golgi apparatus and endoplasmic reticulum (ER). **(d)** Detection of dLight sensor (scattered dark material, green arrowheads) in a dendrite (green outline) establishing synapses (blue arrows) with a TH-positive axon (gold particles, red arrowheads).

### dLight measures both slow and fast/phasic dopamine changes in DLS with *in vivo* pharmacological manipulations

To determine the utility of dLight for measuring dopamine dynamics in DLS *in vivo*, we expressed the sensor in this striatal subregion and measured the fluorescence intensity profile using a custom-built *in vivo* fiber photometry system based on TCSPC principles (**Figure 4a**). The *in vivo* fiber photometry measurement is similar to that used in our previous work (Cui et al., 2013; Cui et al., 2014; Kupferschmidt et al., 2017). We measured dLight fluorescence from the DLS of mice treated with cocaine (15 mg/kg i.p.) after a 10 min observation period in their home cage. We found that cocaine produced an increase in fluorescence that plateaued after 20 min (**Figures 4b & c**). We also observed transient (subsecond to second duration) increases in fluorescence both before and after cocaine administration (**Figure 4d**). To examine the profiles of these spontaneous transients, we time-locked the transients to their peaks (**Figure 4e** **& f**). We then analyzed cocaine-induced changes in transient frequency, amplitude, and decay (**Figure 4g-j**).

**Figure 4.**
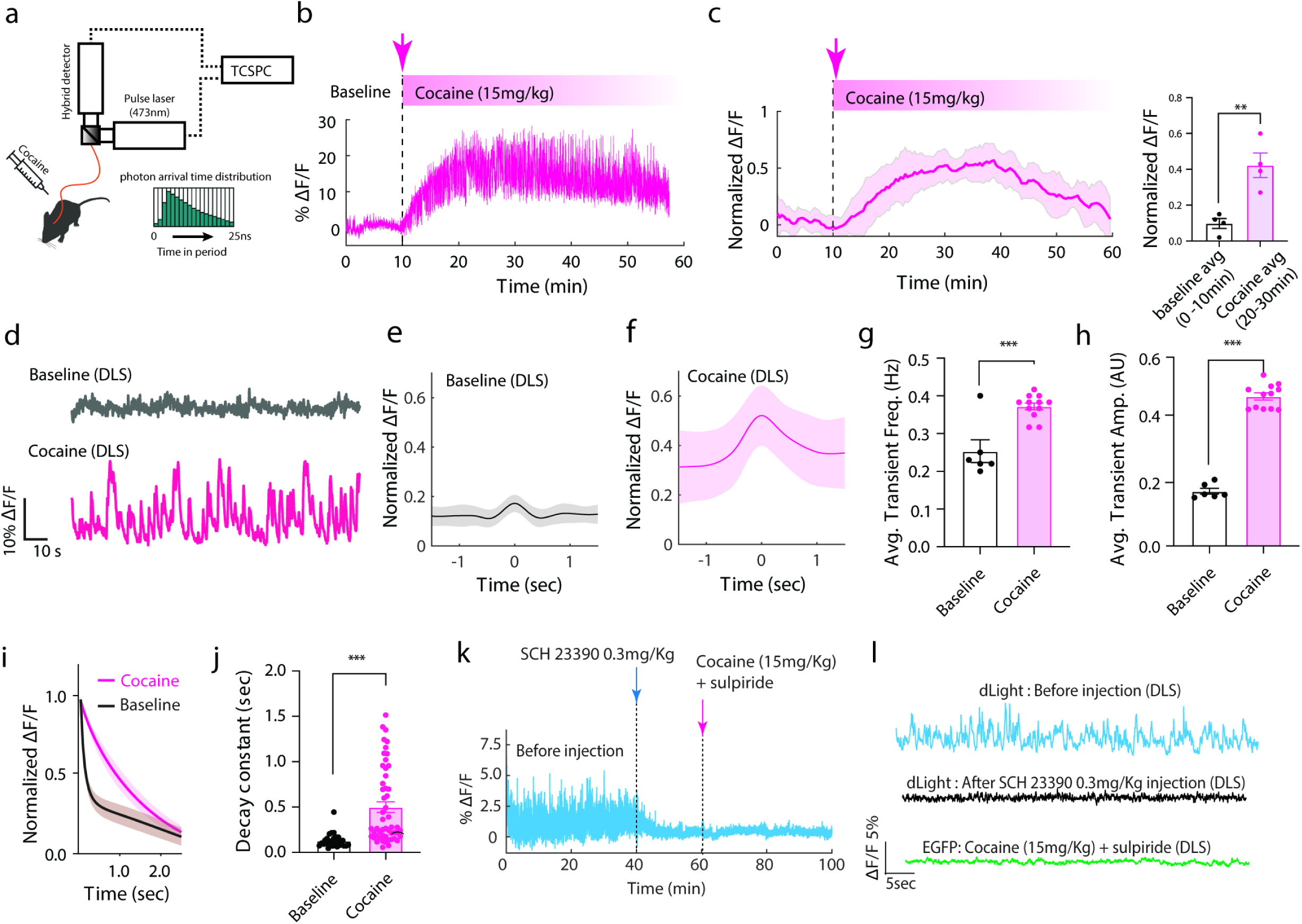
*In vivo* DA measurement in DLS following cocaine administration. **(a)** Schematic diagram of *in vivo* fiber photometry system. **(b)** Sample fluorescence (dF/F_0_, % baseline) profile of DA activity in DLS before and after i.p. cocaine injection. **(c)** Average fluorescence (dF/ F_0_, normalized to session maximum) profile of mice (n=4) before and after i.p. cocaine injection (left panel), and comparison of the average dF/ F_0_ between 10min of baseline and 10 min of cocaine injection (right panel, two-tailed paired t-test ** p=0.0050) **(d)** Magnified dF/ F_0_ profile illustrating DA activity before (Baseline) and after i.p. cocaine injection. **(e)** Averaged fluorescence (dF/ F_0_, normalized to session maximum) profile of baseline DA transients, time-locked to the transient peaks (n=64). **(f)** Averaged fluorescence (dF/ F_0_, normalized to session maximum) profile of DA transients following cocaine injection, time-locked to the transient peaks (n=165). DA transient frequency **(g)** and amplitude **(h)** were increased following cocaine injection. Each point denotes a five minute average. **(i)** Average exponential fit of normalized dopamine transient decay. **(j)** Fitted exponential decay constant values showing longer DA transient decay time following cocaine injection; individual points correspond to individual transients. **(k)** dLight fluorescence is blocked by *in vivo* administration of the D1R antagonist, SCH23390. **(l)** Representative dF/ F_0_ profiles for dLight in DLS before and after SCH23390 administration as well as for an eGFP control mouse. *** p<0.001. Shaded areas and error bars represent the SEM.

We found that cocaine increased dopamine transient frequency (t=4.887, p<0.001, df=16) and amplitude (t=13.81, p<0.001, df=16). Furthermore, we found that cocaine increased the decay time constant of spontaneous dopamine transients (t=4.730 p<0.001, df=79).

### In vivo changes in fluorescence originate from dLight

To determine if the fluorescence changes observed during our recordings originated from dLight and to ensure that the observed changes in fluorescence were due to dopamine actions on dLight (and not altered fluorescence readout due to changes in blood flow, for example), we further analyzed the fluorescence signals in eGFP-expressing mice (**Supplementary Figure 3**) in freely moving conditions and after cocaine injection. In these recordings we did not observe spontaneous, fast transients either before or after cocaine injection. There was also no slow increase in eGFP fluorescence after cocaine injection. We also validated, through benchtop testing, that we do not observe transient changes in fluorescence due to fiber bending or movement artifact when recording with the TCSPC fiber photometry system (**Supplementary Figure 3**). We also performed recordings in which dLight-expressing mice were treated with the D1 dopamine receptor antagonist SCH23390 to block dopamine binding to the sensor. Indeed, treatment with SCH23390 eliminated phasic dopamine transients and this did not change with application of cocaine plus a D2 receptor antagonist (**Figure 4k**). Further, in the presence of SCH23390, the fluorescence profile for dLight resembled that of a static eGFP control mouse (**Figure 4l**).

### dLight measures distinctive phasic dopamine dynamics in DLS in comparison to other striatal subregions during Pavlovian conditioning

Next, we characterized the phasic DA profiles in the DLS using *in vivo* dLight photometry, and assessed regional differences in dopamine transients during Pavlovian conditioning as this training paradigm produces clear behavioral changes associated with consistent and reliable phasic DA release in NAc. We trained animals in a well-established conditioning paradigm for a food reinforcer using a tone as the conditioned stimulus (**Figure 5a-d**). During 14 sessions of daily training, the latency between reward delivery and head-entry decreased and stabilized (**Figure 5b and c**; two-tailed paired t-test, p<0.0001). **Figure 5e** shows example traces of phasic dopamine transients in the *in vivo* dLight photometry measurement across striatal subregions.

**Figure 5.**
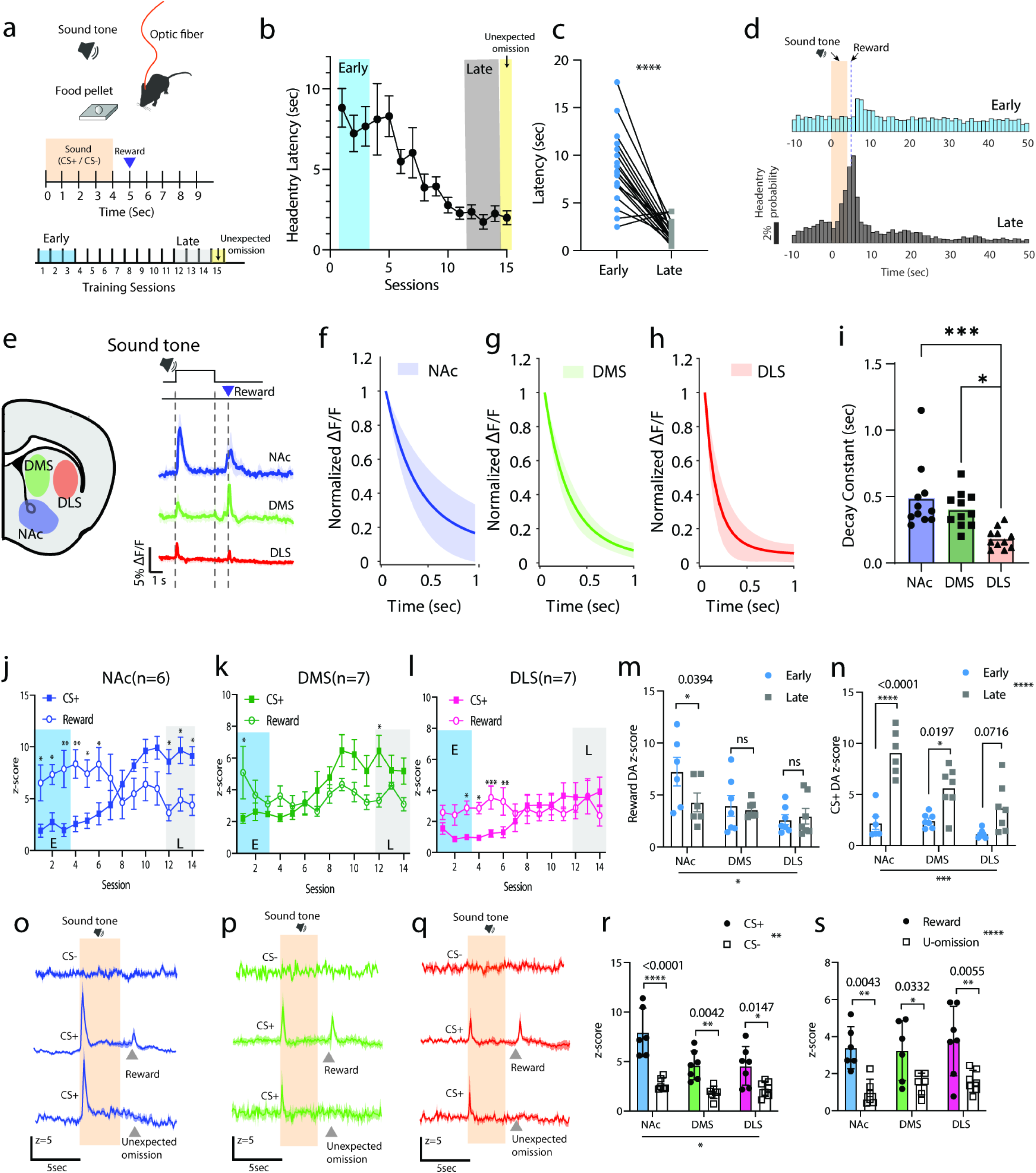
*In vivo* DA dynamics across striatal regions during Pavlovian training. **(a)** Schematic diagram of the experimental design and **(b,c)** behavioral learning curve during Pavlovian training (n=20 mice, two-tailed paired t-test, p<0.0001). **(d)** Head-entry probability in the early (days 1-3) and late training sessions (days 12-14). **(e)** Monitoring sites in the striatum and example traces of dF/ F_0_ session average profiles in NAc, DMS, and DLS. **(f,g,h)** Averaged decay constant fitting of normalized DA transients aligned to the onset of CS+. **(i)** Summarized decay constants across striatal regions. Each point is an average of 15 CS+ trials per animal (one-way ANOVA with Bonferroni’s correction for multiple comparisons, * p=0.01, *** p=0.0005). **(j,k,l)** DA responses to CS+ (open circles) and reward-delivery (filled squares) during the Pavlovian training in NAc (left, blue, n=6 mice two-way ANOVA time x agent interaction: p<0.0001 ****), DMS (mid, green, n=7 mice two-way ANOVA time x agent interaction: p<0.0006 ***), and DLS (right, red, n=7 mice, two way ANOVA time x agent interaction: p<0.0001 ****), respectively. **(m)** DA responses coupled to reward delivery (two-way ANOVA main effect : training : n.s region : p=0.0202). **(n)** DA responses coupled to CS+ in the early and late training sessions (two-way ANOVA region effect : p=0.0202*, training effect : n.s. p=0.1078, region x training interaction: n.s. p=0.0911 F (2, 17) = 2.767) **(o,p,q)** Across striatal regions, DA responses to CS- presentation (top row), CS+ (mid row) and unexpected omission (top row) in the late training phase. **(r)** DA responses coupled to CS+ or CS- presentation (two-way ANOVA, cue effect: p<0.0001**** region effect: p=0.0075 ** cue x region interaction: p=0.0268*). **(s)** DA responses in unexpected-omission test (two-way ANOVA reward effect : p<0.0001**** region : ns p=0.6399, reward x region interaction : ns p=0.7884). Shaded orange areas indicate CS duration.

The average dopamine transients evoked by the CS+ were compared across the striatal subregions (**Figure 5f-h**). We found that the average decay time of individual transients in DLS was significantly faster than in NAc (**Figure 5f-h** **and i**; one-way ANOVA, F(2,27)=14.93, p<0.0001).

A training-dependent temporal shift in the dopamine increases is shown in **Figure 5j,k** **and I** (as well as Supplementary Figure 4 showing a color plot of increases on all training trials). Similar to previous work (Brown et al., 2011; Willuhn et al., 2012; Patriarchi et al., 2018), we observed dopamine transients in NAc with a short latency after reinforcer delivery that gradually shifted to a short latency after the presentation of the CS+ during the conditioning (**Figure 5j,m,n**). The response in NAc associated with reward delivery decreased over the course or training. Interestingly, in the DMS and DLS, the transient response to the reward delivery was maintained late in conditioning (**Figure 5 k,l**), and thus the reward-related DA response was not significantly different across the training period in either DMS or DLS (**Figure 5m**).

Dopamine transients tied to the CS+ showed regionally distinctive occurrence during training (**Figure 5n**; two-way ANOVA, repeated measure, region effect: p=0.0001***, training effect: p<0.0001****, region x training interaction: p=0.0240 *, F(2,17)=4.682). For example, in the NAc and DMS there was a gradual and significant increase in the dopamine transient from the early to late training phase (**Figure 5n**; Sidak’s multiple comparison test, NAc p<0.0001, DMS p=0.0197). Some animals exhibited a CS+-associated dopamine transient during the first few conditioning trials (**Supplementary Figure 4**) presumably in response to the presentation of this novel environmental event. On subsequent trials early in training this response disappeared, but re-emerged later in training.

**Figure 5o-s** **and** **Figure6** show conditioning-related dopamine transients in all three striatal subregions. The reward-related DA transients (left top panels of **Figure 6a-c**) were reduced or absent in unexpected reward omission tests (middle top panels of **Figure 6a-c** **and** **Figure 5s**), despite vigorous head entry activity (middle bottom panels of **Figure 6a-c**, omission rate <17% within a session). Thus, the reward-induced dopamine increases are not simply the product of increased reward-directed behaviors. As expected, in the CS- trials, we observed a lack of dopamine transients accompanied by low levels of head entry activity (right top/bottom panels of **Figure 6a-c** **and** **Figure 5r**). It is important to note that no responses were observed in eGFP-expressing control animals under any of the conditioned stimulus or reward delivery conditions despite the same training and reward-retrieval behavior (**Figure 6d**). Thus, the conditioning related transients observed with dLight are not due to movement of an optic patch cable or other

**Figure 6.**
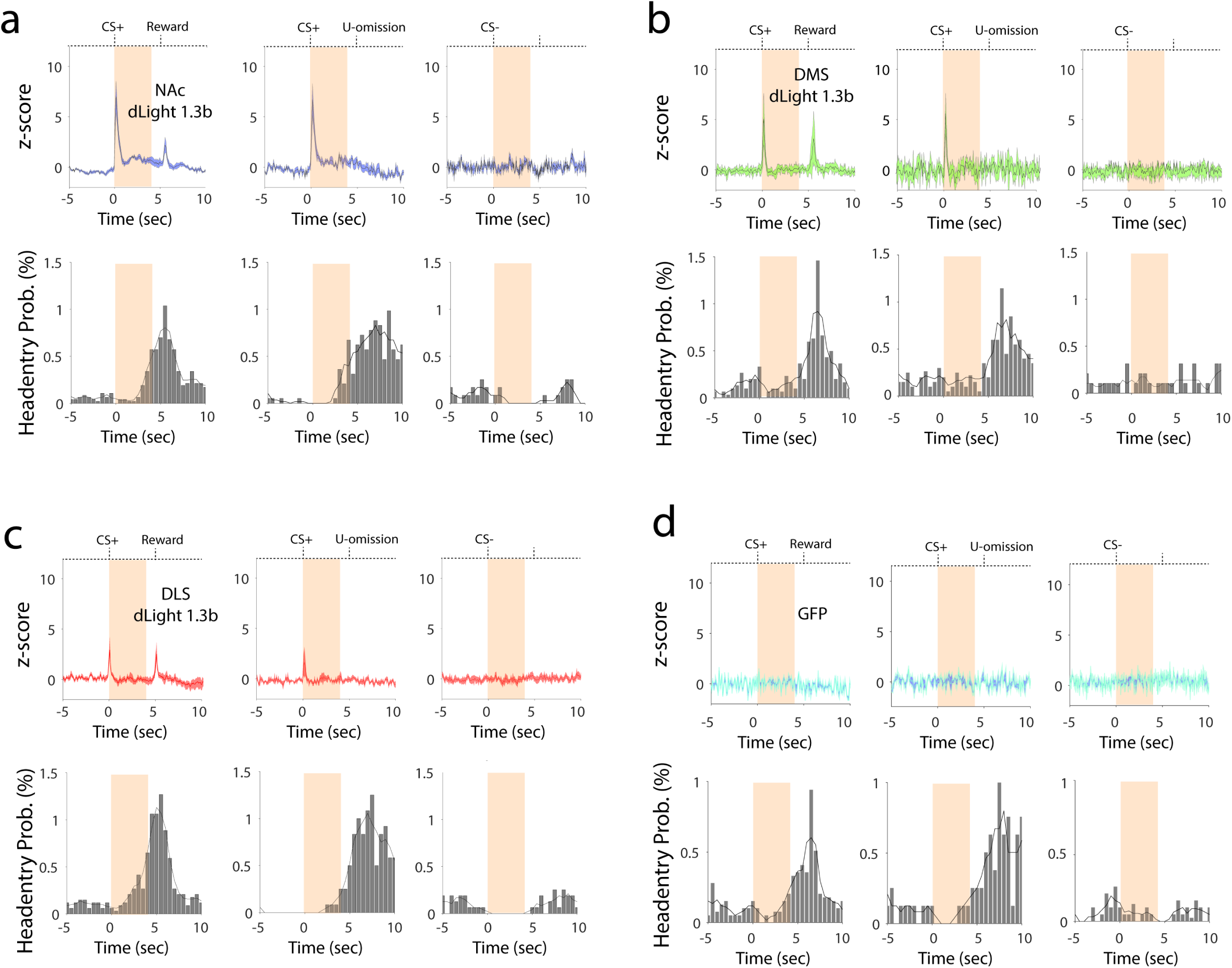
Head-entry rate and DA activity patterns in: **(a)** NAc with CS+ and reward (left) CS+ and unexpected omission (mid), and CS- without reward (right); **(b)** DMS with CS+ and reward (left) CS+ and unexpected omission (mid), and CS- without reward (right); and **(c)** DLS with CS+ and reward (left) CS+ and unexpected omission (mid), and CS- without reward (right), and **(d)** Fluorescence signals in GFP control mice with CS+ and reward (left) CS+ and unexpected omission (mid), and CS- without reward (right). Shaded orange areas indicate CS duration. Note the sustained increase in fluorescence between the offset of the transient induced by the CS+ and reward delivery or unexpected omission in the NAc but not DMS or DLS.

non-physiological changes, but rather appear to reflect changes in dopamine related to conditioning and behavior. Interestingly, we observed sustained fluorescence increases in NAc between 0-5 sec after CS+, but not CS-, presentation regardless of reward delivery (**Figure 5o** **and** **Figure 6a**). These sustained responses were not observed in DMS or DLS (**Figure 5p,q**, **Figure 6b,c**; **Supplementary Figure 5**).

## Discussion

We first used dLight photometry to examine electrical stimulation-induced dopamine increases, with simultaneous FSCV in brain slice experiments. In general, we found similar responses with both techniques across a range of stimulus intensities. We did detect photometric signals at slightly lower stimulus intensities, but still within the range where FSCV signals are observed in some studies (e.g. Salinas et al., 2016). We also found that the dLight signal was near maximal at mid and higher stimulation intensities (>400 µA) sometimes used in FSCV. Several observations indicated that the electrical stimulation-induced changes in fluorescence we observed were indeed due to dLight activation by dopamine released from nigrostriatal afferents. The transients were blocked by a D1R antagonist, strongly reduced following virally expressed Caspase3 ablation of DAT Cre+ substantia nigra neurons, and no responses were observed in slices expressing eGFP in place of dLight. We next sought to determine if regulation of dopamine release by the D2 autoreceptor would be similar when comparing the two methods and found that application of quinpirole (D2 dopamine receptor agonist) resulted in a similar average inhibition of evoked dopamine release measured with both techniques.

Interestingly, evoked photometric responses were nearly maximal at stimulation intensities yielding a ∼50% maximal voltammetric response. It is difficult to compare the sensitivity of the two techniques, as one needs to account for differences in sampling volumes that yield a response value per unit of measurement (e.g. 5% dF/ F_0_ or 500nM DA per µm^3^). Thus, the fundamentally different sampling volumes for each method preclude statements about direct differences in sensitivity. Nonetheless, the inherently greater sampling volumes in *in vivo* and *in vitro* photometric methods facilitate the detection of dopamine (or any “volume” neurotransmitter) in a way that is not possible with single electrode voltammetric methods. We must emphasize, however, that our findings do not indicate that dLight is a more sensitive technique for detecting dopamine release. Findings with the two techniques cannot be compared directly, given factors including differences in the locations of dopamine detection and the carbon fiber dimensions.

However, it seems safe to say that dLight photometry has a detection capability comparable to FSCV.

### Slow and fast/phasic dopamine dynamics

Striatal (and likely extra-striatal) dopamine operates on multiple time scales and dopamine levels, including tonic as well as slow and fast/phasic changes (Liu et al., 2021). Tonic dopamine levels are set by the basal firing rate of midbrain dopamine neurons and possibly modulated by local striatal mechanisms. Slowly-developing and sustained changes in dopamine occur, e.g. in response to application of DAT blockers. In contrast, phasic dopamine changes are faster, typically lasting on the order of seconds, and are mediated by burst firing of midbrain dopamine neurons, local control of dopamine release (e.g. by acetylcholine or glutamate receptors on dopamine axons), or both (Zhou et al., 2001; Cachope et al., 2012; Threlfell et al., 2012). Slow and phasic dopamine play distinct roles in behavior and motivation (Beeler et al., 2010; Hamid et al., 2016; Berke, 2018; Mohebi et al., 2019; Wang et al., 2021). Microdialysis sampling with electrochemical detection techniques allows for measurement of absolute concentrations of dopamine that reach the probe (at least when using variants such as no-net-flux). Assessment of tonic dopamine levels within and between test sessions can also be obtained with microdialysis, but cannot be used to assess phasic changes in dopamine levels on a behaviorally-relevant time scale.

Like FSCV, dLight photometry cannot measure the absolute tonic extracellular dopamine concentration. The subsecond sampling in FSCV allows for measurement of phasic dopamine levels but due to the need to subtract a recent baseline signal, it is generally not useful for determining slow changes in dopamine levels. A variant of FSCV using an altered waveform (known as FSCAV; Burrell et al., 2015) can be used to measure slower changes in dopamine, but requires slower data acquisition (10s or sec) that precludes simultaneous measurement of phasic dopamine changes. Given these limitations, dLight (and other fluorescent biosensors) represents a technological advance. As we demonstrate here, dLight allows for simultaneous assessment of slow and phasic dopamine changes within an *in vivo* testing session. This was evident in our experiments examining the effect of acute cocaine treatment on DLS dopamine levels where we observed an increase in baseline dopamine levels that developed over several minutes following cocaine administration. This was accompanied by increased amplitude, frequency, and duration of phasic dopamine transients. In contrast to our *in vitro* work, we observed a cocaine-induced increase in transient amplitude *in vivo*. This is likely due to the slower sampling rates used for our *in vivo* recordings (20 Hz) which may slightly underestimate phasic responses to the highest burst firing frequencies of dopamine neurons (up to 100Hz) which are thought to underlie phasic dopamine transients (Hyland et al., 2002; Lohani et al., 2018). Indirect cocaine effects may also contribute to increased transient amplitudes *in vivo*. For example, cocaine could alter midbrain dopaminergic neuron firing *in vivo*, and this could also contribute to transient amplitudes (Covey et al., 2014). The dopamine transients in DLS preferentially reflect locomotion in naturalistic motor behaviors (Markowitz et al. 2023) and even after cocaine injections (Jørgensen et al. 2023). Increased locomotion induced by cocaine injection may also contribute to increases in transient amplitude and frequency. Such effects would not contribute to dopamine release in brain slices.

An understanding of the significance of slow and fast/phasic dopamine changes is important. Changes in phasic dopamine release can occur in the absence of slower changes in dopamine levels and vice versa. Also, tonic and slow, long-lasting changes in dopamine levels may serve to influence global striatal activity or metaplasticity. For example, the chronic DA deficiency observed in PD models is often accompanied by changes in synaptic plasticity or altered cellular physiology (Calabresi et al., 2009; Surmeier et al., 2014; Wang and Zhang, 2016; Zhai et al., 2018; Graves and Surmeier, 2019). Thus, the ability to simultaneously assess slow and phasic changes in dopamine release in freely behaving animals will be of great use to the field.

### Technical considerations for sampling

Sampling rates can be much faster with photometry than with FSCV. FSCV sampling is inherently limited by the electrochemical properties of the triangle wave form (∼8ms) and the time required for desorption of dopamine from the CFE surface, resulting in a maximum practical sampling rate of up to ∼50Hz (though 10Hz is typical for dopamine measurements). In contrast, the sampling rates for photometric methods are limited by the binding and unbinding rates of the sensor and the digitization rate of the data acquisition hardware used. Our *in vitro* photometric responses were collected at 100Hz or 1,000Hz, although rates up to 20,000Hz are possible with our hardware configuration. *In vivo* fiber photometry sampling rates with most available systems can also exceed 100Hz. However, given the known time course of dopamine signaling and dLight kinetics we chose to perform our recordings at 20Hz. It is important to note that, with photometric methods, the fluorescent on/off rates of dLight (or other biosensors) will limit the utility of high sampling rate data. For example, the on and off rates for dLight1.1 are ∼10 and 100ms, respectively (Patriarchi et al., 2018; Sabatini and Tian, 2020). Thus, it is possible that sampling at 200Hz with dLight1.1 will yield a large data set that would not differ practically from a data set collected at 100 or even 20Hz. These fluorescence kinetics may also limit interpretation of transient rise times or modeling of decay kinetics, as the fluorescent signal may not actually represent the termination of neuromodulator signaling but rather the off rate of the sensor. Thus, consideration of the experimental question and the type of data that can be collected should be considered when choosing a fluorescent biosensor (Labouesse et al., 2020; Sabatini and Tian, 2020).

### Synaptic dopamine vs overflow

Using dLight it may be possible to assess truly “synaptic” dopamine release. That is to say, both microdialysis and FSCV are methods that measure dopamine overflow out of the synapse collected or detected at a site distant from most release sites. Because of the size of the probes, sampling of dopamine at synaptic release sites has not been possible with either of these methods. In this context, dLight represents another advance for the field because it can be genetically targeted to distinct cellular compartments. For example, dLight sensors are integral membrane proteins so they will only be expressed in membranes. We confirmed this with our electron microscopy work and show that indeed, dLight traffics strongly to plasma membrane, Golgi, and ER compartments of the neuron (**Figure 3c**). Further, our results show that dLight traffics to synaptic and nearby extrasynaptic sites (**Figure 3b,d**). Thus, dLight signals likely represent dopamine release that acts at these proximal sites. Work from the Ford and Williams labs (Ford et al., 2009; Marcott et al., 2014; Mamaligas et al., 2016) utilized an indirect electrophysiological approach to measure GIRK-mediated currents activated by D2 dopamine receptors. Like the dLight method we used, this approach likely represents a mixture of synaptic and extrasynaptic actions of dopamine, but is not amenable to *in vivo* use. More recent work from the Williams lab used two photon excitation with dLight to measure dopamine release at spatially discrete sites in the midbrain (Condon et al., 2021). These authors concluded that the time course of D2 receptor-mediated responses was dictated largely by dopamine release and not diffusion. Given the sparsity of dopamine release sites in midbrain, it is possible that these spatially discrete sites represent dopamine synapses. Thus, the use of dLight with high resolution imaging methods may allow for measurement of synaptic dopamine release. Furthermore, with dual color fluorescence imaging methods, synaptic markers could be employed to allow for labeling of pre or post-synaptic elements to colocalize with dLight (or other biosensor) signals, facilitating measurement of truly synaptic neuromodulator release.

### A new method settles an old controversy

Pharmacologically, the direct mechanism of action of cocaine is inhibition of DAT (and other monoamine transporters). This should result in prolonged phasic dopamine transients and, in regions where the DAT function is the primary mechanism of dopamine clearance, increases in tonic dopamine levels. Indeed, with microdialysis, the expected increases in tonic dopamine have been confirmed (Church et al., 1987; Hurd et al., 1988). Cocaine increases the duration of evoked dopamine transients measured with FSCV. Interestingly, measurements with FSCV show that cocaine (and other DAT blockers) also increase the observed dopamine transient peak amplitude in brain slices. This is often interpreted to mean that DAT blockers enhance or facilitate electrically evoked dopamine release in slice. From a pharmacological perspective, this is not intuitive as dopamine release should not be affected by DAT inhibition. Interestingly, Patriarchi et al. (2018) noted that there was no increase in dLight dopamine transient peak amplitude in response to cocaine application. Following up on this, we examined cocaine effects on dopamine transient peak amplitude and duration in our simultaneous dLight and FSCV recordings. Similar to previous work, we observed a large cocaine-induced increase in dopamine transient peak amplitude with FSCV but only modest or no effects with dLight (**Figure 2**). With both methods, the dopamine transient decay was prolonged by the drug. Similar results were obtained with the more specific DAT blocker nomifensine. To examine this discrepancy, we first considered that the physical interaction of dopamine with the carbon fiber electrode may be the culprit. We thus modified the triangle waveform used in typical FSCV to reduce dopamine adsorption/sensitivity at the carbon fiber electrode. This modification resulted in a loss of the nomifensine-induced increases in dopamine transient peak amplitude normally observed in FSCV recordings with the traditional voltage triangle waveform (**Figure 2h-j**). We posit that if DAT inhibitors did in fact increase dopamine release (Wu et al., 2001; Venton et al., 2006; Oleson et al., 2009; Kile et al., 2010; Yorgason et al., 2011; Covey et al., 2014; Hoffman et al., 2016; Salinas et al., 2016), then the change in CFE sensitivity to dopamine with the modified waveform should not adversely affect putative DAT blocker-induced increases in dopamine release.

In a follow up experiment, we applied cocaine to dorsal striatal brain slices from DAT KO mice. We did not observe any cocaine-induced increase in dopamine release (**Figure 2**) in DAT KO mice. Our observation that dopamine transients were not prolonged by cocaine in DAT KO mice is consistent with the lack of functional transporter. These findings are in line with previous work in DAT KO mice showing no contribution of other catecholamine transporters to DA clearance in NAc (Budygin et al., 2002; Mateo et al., 2004).

Altogether, our data do not support the idea of DAT blocker-induced increases in dopamine release. Therefore, we posit the following alternative interpretation of the observed DAT blocker-induced increases in dopamine transient peaks using FSCV. Rather than increasing or facilitating dopamine release, DAT blockers by virtue of their inhibition of dopamine clearance allow for an increase in the point diffusion of dopamine from its release site. That is to say, under normal conditions, the diffusion of dopamine is limited by several factors, most notably DAT (Cragg and Rice, 2004; Hoffman et al., 2016), but under conditions of impaired DAT function (e.g. in the presence of a DAT blocker) the diffusional spread of dopamine from its release site is increased (Venton et al., 2003). This would effectively increase the sampling volume of CFEs allowing for greater FSCV detection of dopamine (and increased dopamine transient peak heights) in the absence of an actual increase in release. Further, because the sampling volume used in photometric methods is already much larger than with FSCV methods, the cocaine induced increase in dopamine diffusion from its release site has a more negligible effect on dLight/photometric DA transients. Thus, we believe that dLight more faithfully reflects dopamine release spatial dynamics than FSCV, especially under conditions where dopamine overflow or uptake may be affected.

### dLight allows for insights into striatal subregion dopamine release differences *in vivo*

We compared dopamine transients across the three striatal subregions: the NAc, DMS, and DLS as animals learned a Pavlovian association between a CS+ and reward. We found that dopamine transient decay times were faster in DLS than in NAc, with intermediate levels in DMS. These findings are consistent with the kinetics of electrical stimulation-induced dopamine transients measured in brain slices and *in vivo* (Garris et al., 1994; Calipari et al., 2012; although see Brown et al, 2011). However, there is to date little information about subregional differences in dopamine transients driven by environmental stimuli or associated with behavior and measured *in vivo*.

Our findings reveal several interesting differences in dopamine dynamics over the course of Pavlovian conditioning in the different striatal subregions. While responses to the CS+ generally increase over the course of training, the responses to reward delivery are maintained in DMS and DLS, but reduced over training in NAc. These responses in dorsal striatum did not track with head entries, as observed in the unexpected omission trials (Figures 5p,q and 6b,c), indicating that they are dependent on reward delivery per se. The CS+ and reward-related responses in NAc are consistent with previous findings examining dopaminergic neuronal firing and FSCV (e.g. Roitman et al., 2004; Schultz et al., 1997). The sustained elevation between CS+ delivery and reward is not always observed, but has been seen in past studies that used FSCV or dLight fiber photometry during performance of *in vivo* learning tasks (Hamid et al., 2016; Mohebi et al., 2019). In our experiments this increase appeared to be tied to expectation of reward delivery (i.e. Figures 5o, 6a and Supplementary Figure 4). Sustained responses of this type were not observed in DLS or DMS. It will be interesting to determine the neural basis of this sustained increase in dopamine, as well as if and how it contributes to task performance.

*In vivo* fiber photometry technical difficulties include potential artifacts related to fiber bending which can result in light exposure to fiber cladding, usually related to animal movement. Thus, it is important to include crucial analyses and control procedures that can detect such contaminating signals. We and others have used time-correlated single photon counting-based fiber photometry to measure signals with a variety of genetically-encoded sensors in tasks including the open field, operant lever pressing and the accelerating rotarod (Cui et al., 2013; Cui et al., 2014; Kupferschmidt et al., 2017). In general, we find few artifacts due to light entering the cladding unless fibers are severely kinked which generally does not happen in open field or operant box settings. Unlike continuous-wave (CW) laser or light emitting diode (LED), the MHz pulsed laser system used in the TCPSC measurement also minimizes the period of light exposure to the fiber cladding and has a high-pass filtering effect that reduces the likelihood of fiber bending based artifacts. This system does not include the “isosbestic” excitation control used in several fiber photometry systems (Kim et al., 2016). Instead, using eGFP alone fluorescent controls in the current study and in previous work from the lab we observed no evidence of artifactual changes in fluorescence (Cui et al., 2013; Kupferschmidt et al., 2017). For these reasons we are confident that the dLight signals we measured *in vivo* with TCSPC truly indicate changes in striatal dopamine. Likewise, several control experiments including recordings with eGFP alone, blockade of the signal by SCH23390 and loss of signal after lesioning SNc neurons indicate that the stimulus-induced fluorescence increases we observed in brain slices reflect dopamine increases, and these findings are consistent with data from our previous studies indicating that photometry signals using this approach are not contaminated by endogenous fluorescence or other non sensor sources (Sgobio et al., 2014; Kupferschmidt and Lovinger, 2015).

Dopamine release in DLS was measured previously using FSCV in rat conditioning and y-maze paradigms (Brown et al. 2011, Howe et al., 2013, Klanker et al. 2017). The findings in NAc and DMS in the Brown et al. study are similar to those that we observe with dLight fiber photometry in well-trained mice. However, measurements with FSCV in DLS show little-to-no fast dopamine release in response to unexpected food reward or a positive discriminative stimulus (DS+), and a non-significant slower-developing increase that persists for a few seconds after DS+ presentation. The behavioral correlate of this late, slow component of dopamine release is unclear, but it might be related to movement. From this study it was not clear if the lack of dopamine changes in DLS were due to the inability of FSCV to detect dopamine in this region or if DLS dopamine increases are not produced by reward or predictive stimuli. Our findings indicate that both stimulus and reward-related dopamine increases can be detected in DLS in mouse using dLight fiber photometry, consistent the former conclusion. It should also be noted that Brown and coworkers did not provide information regarding the time course of dopamine release changes over the course of training, as we have been able to do with dLight fiber photometry. In the Howe et al. (2013) study, gradual “ramping” increases in dopamine were observed in DLS on less than half the trials as rats approached the goal in a Y-maze task, but no fast/phasic changes in dopamine were observed. The Klanker et al. (2017) study showed small increases in DLS that appeared to be related to movement initiation in a well-learned operant task, while van Elzelingen et al. (2022) showed dopamine signals in rat DLS during Pavlovian conditioning behavior. These responses were not followed over the course of initial training, but were examined during the course of reversal training. Willuhn and coworkers (2012) were able to detect fast/phasic dopamine increases in rat DLS that developed over the course of cocaine self-administration using FSCV. However, the prolongation of dopamine increases produced by this DAT blocker likely facilitated detection of this increase. It should now be clear that past studies showed dopamine detection in rat DLS, but our findings provide new information on responses to drugs and environmental events in this important striatal subregion in mouse, as well as evidence that we can measure both slow and fast changes in dopamine simultaneously throughout the course of pharmacological and extended behavioral studies.

## Materials and Methods

### Subjects

Three-month-old, male C57BL/6J mice were obtained from the Jackson Laboratory (Strain 000664) and pair-housed in the vivarium for at least one week before any experimental use. DAT-IRES-Cre mice were obtained from the Jackson Laboratory (Strain 006660) and bred in house. DAT KO mice were developed as previously described (Giros et al., 1996) and obtained from the Sara Jones laboratory at Wake Forest University. Male and female DAT-IRES-Cre and DAT KO transgenic mice were used in all experiments. All mice were housed with 2-4 mice per cage and maintained on a 12:12 hour light cycle and ad libitum access to food and water.

All procedures performed in this work follow the guidelines of the Institutional Animal Care and Use Committee of the Division of Intramural Clinical and Biological Research, National Institute on Alcohol Abuse and Alcoholism, and the animal care and use protocol was approved by this committee.

### Viruses and Stereotaxic Injections

The Caspase3-coding virus (AAV1-EF1a-FLEX-taCasp3-TEVp) was custom packaged by Vigene Biosciences and a gift from Dr. Huaibin Cai, National Institute on Aging. dLight1.1 viruses were used for *in vitro* experiments and were either generated in the Tian laboratory and sent to NIAAA (AAV5-CAG-dLight1.1 and AAV9-CAG-dLight1.1) or purchased from Addgene (pAAV5-CAG- dLight1.1; Addgene viral prep # 111067-AAV5). AAV9-CAG-dLight1.1 are only partially used in Fig2. All other in-vitro experiments used AAV5-CAG-dLight1.1. dLight1.3b virus was used for *in vivo* experiments and supplied by theTian laboratory or custom packaged by Vigene Biosciences (AAV9-CAG-dLight1.3b).

All stereotaxic injections were conducted using sterile technique on mice at least 3 months of age. Mice were anesthetized with a 5% isoflurane/oxygen mixture and placed in a Kopf stereotaxic frame. Anesthesia was maintained with 1-2% isoflurane/oxygen mixture. The skulls were leveled and an incision was made to expose the skull. Craniotomies were made over the dorsal striatum (AP +1.0, ML +/-1.8, from Bregma in mm), nucleus accumbens (AP +1.2, ML +/-0.8, from Bregma in mm) or substantia nigra (AP -3.0, ML +/-1.2, from Bregma in mm) and a 1 µL Neuros Hamilton Syringe was lowered slowly to the desired depth from the brain surface (-2.25mm, -3.75mm, and -4.1mm for dorsal striatum, nucleus accumbens, and substantia nigra, respectively). For in-vivo experiments, viruses were injected in the following coordinates. NAc (AP +1.2, ML +/-0.8, DV - 3.75), DMS (AP +1.0, ML +/-1.2, DV -2.25), DLS (AP +0.8, ML +/-2.2, DV -2.25) where AP and ML in mm from Bregma, DV in mm from the brain surface. Virus infusion volumes were 300nL for dorsal striatum and 500nL for VTA and viruses were infused at 50nL/minute. After infusions were completed, the syringe was left in place for 10 minutes before withdrawal and the incisions were closed with VetBond. Mice recovered for at least three weeks before being used for in vitro experiments or fiber optic implantation for *in vivo* experiments.

### Simultaneous Photometry and Voltammetric Recordings

Dual photometric and voltammetric recordings were conducted as previously described (Sgobio et al., 2014; Sgobio et al., 2019) except that the photometry recordings in the current study used dLight1.1 or dLight1.3b to compare dopamine release between voltammetric and photometric methods simultaneously in the same preparation. Brain slices containing the striatum were prepared as previously described (Salinas et al., 2016). Briefly, mice were anesthetized with isoflurane, rapidly decapitated, brains extracted, and 300µm thick coronal sections prepared on a vibratome (Leica VT 1200S). The slices were hemisected and inspected to ensure viral expression of dLight in the region of interest using an epifluorescent Zeiss AxioZoom microscope equipped with a GFP filter set (Carl Zeiss Microscopy Filter Set Lumar # 38 BP470/40, FT495, BP525/50). Then the slices were incubated at 32°C for 30 minutes before being moved to room temperature for one hour before beginning experiments.

Brain slices with dLight expression were moved to an upright Zeiss AxioSkop2 microscope mounted on a XY translational stage and equipped with a GFP filter set. Oxygenated ACSF was perfused at 1.5-2mL/minute and warmed to 30-32°C. The recording region of interest was located under 4X magnification and fluorescent illumination to ensure dLight expression in the region of interest. Then a stainless steel twisted bipolar stimulating electrode (P1 Technologies) was placed on the tissue surface near the area of dLight expression. For simultaneous photometry voltammetry recordings, carbon fiber electrodes (CFE) were fabricated as previously described (Salinas et al., 2016; Salinas et al., 2021) and placed in the tissue in the center of the recording region of interest under 4X magnification. Slices were then visualized with 40X objective (0.8 NA) and the field of view (∼180um x 180um) was adjusted so the stimulating electrode was just outside the field of view and the CFE was in the center of the field of view. Under 40X magnification, the focus was adjusted to a focal layer beneath the slice surface where fluorescent cells could be identified. Fluorescent transients were quantified with a PMT-based system (PTI D-104 photometer) coupled with a Digidata 1322A (Molecular Devices LLC) to digitize the PMT signal (100-1,000Hz). Clampex software was used to collect photometry data and synchronize photometric and voltammetric recordings through the Digidata. A mechanical shutter (Uniblitz V25) was used to limit exposure to fluorophore-exciting light to discrete periods and minimize photobleaching of the dLight between recordings. FSCV recordings were carried out as previously described (Sgobio et al., 2014; Salinas et al., 2016; Sgobio et al., 2019; Salinas et al., 2021). Briefly, a triangle wave form voltage was applied to the CFE beginning at -0.4V to +1.2V and back to -0.4V. This scan was applied at 400V/s and repeated at 10Hz. Dopamine was identified electrochemically by the oxidation peak at +0.6V on the ascending phase of the triangle ramp. Dopamine release was evoked with electrical stimulation delivered every three minutes using a constant current stimulus isolator (DigiTimer DS3). Input-output (IO) curves were generated to examine evoked dopamine release measured with both techniques across varying electrical stimulation intensities (50-800 µA, 1 ms). For pharmacological experiments, a stimulation intensity yielding approximately 30-60% of the maximal responses with both methods was used to ensure that any subsequent treatments were not limited by floor or ceiling response effects. Baseline responses were collected for 12-20 minutes before drugs (dissolved in ACSF) were bath applied as indicated for each experiment.

### *In Vivo* Fiber Photometry

At least three weeks after virus infusion surgeries, mice used for *in vivo* experiments underwent a second surgery to implant the optical fiber, similar to those previously described (Kupferschmidt et al., 2017). Briefly, mice were anesthetized and mounted into the stereotaxic apparatus and an incision was made in the scalp along the skull midline as before. The skull was cleaned with a 3% H_2_O_2_ solution to remove any connective tissue and the skull was scored several times with a scalpel to create a better surface for the dental cement headcap at the end of the procedure. Craniotomies were made over the brain region of interest and two distal sites for anchor screw placement. Fiber implants (Thorlabs, # CFMC 12L05) were placed in the following coordinates: NAc (AP +1.2, ML +/-0.8, DV -3.75), DMS (AP +1.0, ML +/-1.2, DV -2.25), DLS (AP +0.8, ML +/-2.2, DV -2.25) where AP and ML in mm from Bregma, DV in mm from the brain surface. The anchor screws were placed before proceeding to the fiber optic placement. To ensure optimal fiber optic placement, fluorescence intensity was monitored intraoperatively using a custom designed fiber photometry system consisting of a 473 nm picosecond pulsed laser, a HPM-100- 40 hybrid detector, and a SPC-130EM time correlated single photon counting (TCSPC) module and software (Becker & Hickl). The measurement rate for the fluorescence lifetime and intensity profile was set at 20Hz during all animal experiments. The output was a custom multimode patch cord from Thor labs with a 0.22NA and 200µm fiber core diameter terminating in a 2.5mm ceramic ferrule. The patch cord ferrule was connected to the fiber optic cannula (CFMC22L05, Thor Labs) to be implanted with a ceramic mating sleeve. The implantable fiber optic cannula was lowered slowly into the craniotomy over the brain region of interest while the fluorescence intensity at the desired emission wavelengths (∼500-540 nm) was monitored. The fluorescence intensity typically increased as the fiber optic approached the area of dLight expression. Once fluorescence intensity plateaued, typically ∼300-500µm dorsal to the virus infusion site, the fiber optic cannula was cemented in place and the surgical site was closed around the dental cement headcap. Mice recovered for at least two weeks before further experiments.

For the Pavlovian conditioning experiment, the TCSPC system was synchronized to the operant behavior boxes (MED-PC) through a TTL channel, in which the TTL pulse generated by MED-PC controller was used to trigger the TCSPC system acquisition for 10s prior to each trial (CS+ or CS-) and maintained 60s of data acquisition. The pulsed laser was continuously turned on and sustained stable laser illumination during the entire session. *In vivo* pharmacology measurements were continuously recorded for 60min. Before the recording, the animals were tethered with an optical patch cord with laser illumination, and habituated for 20 min in the open-field arena without recording.

### *In Vitro* Fiber Photometry Benchtop Testing

To test the possibility of patch cable vibration effects on fluorescent signals captured with TCSPC we devised a benchtop testing system (Supplementary Figure 3). The tip of patch cable connect to our *in vivo* photometry system was mounted on an optical table, and a sample tube containing a fluorescent polymer (World Precision Instruments, cat # KWIK-CAST) was placed at the tip of patch cable. Single-photon counting measures of fluorescence using the same laser and detector used for *in vivo* photometry were carried out for 5 min under conditions in which the patch cable was stable, and when the cable was vibrated by movements simulating those that occur was mice traverse the behavioral apparatuses used in our experiments.

### Pavlovian Conditioning

At least three weeks following fiber optic implant surgeries, mild food restriction was carried out to gradually reduce the animals’ body weight to 85-90% of their initial body weight. Thus, chow was limited to 2–3 g per day for 3 days. Concurrently, mice were handled and tethered to an optical patch cord cable within the designated behavior training time. For the first 2 days of pre training, mice received habituation sessions in the operant training box for 1-hr while tethered to the optical patch cord. In the final day of pre-training, mice underwent a magazine training session in which 30 food pellets were delivered on a random-interval schedule (RI-60). No stimulus cue was provided during these sessions. After 3 days of food restriction and pre-training, mice underwent Pavlovian conditioning sessions. The discrimination training entailed two conditioned stimuli; one conditioned stimulus (CS+) was a high-frequency auditory cue followed by the unconditioned stimulus (US, one 15mg food pellet) as a reinforcer, and a second stimulus (CS-) was a low-frequency auditory cue which was not accompanied by a reinforcer delivery. Within a single session, mice received 20 CS+ trials and 20 CS- trials. In the operant conditioning box within sound and light attenuating enclosures, the auditory cue was presented for 4 seconds with random intertrial intervals (60-sec to 120-sec). In the CS+ trials, an US reinforcer was delivered to a pellet receptacle 1 second after the CS+ cue offset. *In vivo* photometry measurements were conducted every other day during the training, but the mice underwent the same continuous daily Pavlovian training sessions while tethered to an optical patch cord (Thor Labs, 200um core, 3 m length) even if photometry measurements were not conducted. Therefore, DLS and DMS mice received 10 weeks of consecutive daily Pavlovian training. NAc mice received four weeks of Pavlovian training. All animals received the behavior training with 24 (+/-2) hours of interval at their own designated time, and animals were acclimated to the behavior room for at least 1hr before all training sessions.

### Fluorescence microscopy

For anatomical studies, dLight virus injected mice were deeply anesthetized with chloral hydrate (35 mg per 100 g), and perfused transcardially with 4% (wt/vol) paraformaldehyde (PFA) with 0.15% (vol/vol) glutaraldehyde and 15% (vol/vol) picric acid in 0.1 M phosphate buffer (PB, pH 7.3). Brains were left in this fixative solution for 2 h at 4 °C, solution was replaced with 2% PFA and left overnight at 4 °C. Brains were rinsed with PB, and cut into coronal serial sections (40 μm thick) with a vibratome (Leica). All animal procedures were approved by the National Institute on Drug Abuse Animal Care and Use Committee.

Free floating coronal vibratome sections were incubated for 1 h in PB supplemented with 4% bovine serum albumin (BSA) and 0.3% Triton X-100. Sections were then incubated with mouse anti-tyrosine hydroxylase (TH) primary antibody (1:1,000 dilution, Millipore-Sigma, Cat# MAB318, RRID: AB_2201528) and guinea pig anti GFP (1:500, Nittobo Medical, Cat# GFP-GP-Af1180, RRID: AB_2571575) overnight at 4°C. After rinsing 3 × 10 min in PB, sections were incubated in Alexa Fluor 594-affiniPure donkey anti-mouse (1:100 dilution, Jackson Immunoresearch Laboratories, Cat# 715-585-151, RRID: AB_2340855) and Alexa Fluor 488-affiniPure Donkey anti guinea pig (1:100, Jackson Immunoresearch Laboratories, Cat# 706-545-148, RRID: AB_2340472) for 2 h at room temperature. After rinsing, sections were mounted on slides and air-dried. Fluorescent images were collected with Zeiss LSM880 Airyscan Confocal System (Zeiss, White Plains, NY). Images were taken with 20× objectives and z-axis stacks were collected at 1 µm. The confocal images were collected from 3 mice.

### Electron microscopy

Vibratome tissue sections were rinsed with PB, incubated with 1% sodium borohydride in PB for 30 min to inactivate free aldehyde groups, rinsed in PB, and then incubated with blocking solution [1% normal goat serum (NGS), 4% BSA in PB supplemented with 0.02% saponin] for 30 min. Sections were then incubated with guinea pig anti-GFP primary antibody (1:500 dilution, Nittobo Medical, Cat# GFP-GP-Af1180, RRID: AB_2571575); or both guinea pig anti-GFP (1:500 dilution) and mouse anti-TH (1:1,000 dilution) for 24 h at 4°C. Sections were rinsed and incubated overnight at 4°C in the secondary antibody goat anti-guinea pig IgG Fab fragment coupled to 1.4 nm gold (1:100 dilution for GFP detection; Nanoprobes Inc., Cat# 2055, RRID: AB_2802149); or the corresponding secondary antibodies: biotinylated goat anti-guinea pig antibody (1:100 dilution for GFP detection; Vector Laboratories, Cat# PK-4007; RRID:AB_2336816) and secondary antibody goat anti-mouse IgG coupled to 1.4 nm gold (1:100 dilution for TH detection; Nanoprobes Inc., Cat# 2001, RRID:AB_2877644). Sections were incubated in avidin-biotinylated horseradish peroxidase complex in PB for 2 h at room temperature and washed. Peroxidase activity was detected with 0.025% 3,3′-diaminobenzidine (DAB) and 0.003% H_2_O_2_ in PB for 5-10 min. Sections were rinsed in PB, and then in double-distilled water, followed by silver enhancement of the gold particles with the Nanoprobe Silver Kit (2012; Nanoprobes Inc., Stony Brook, NY) for 7 min at room temperature. Next, sections were rinsed with PB and fixed with 0.5% osmium tetroxide in PB for 25 min, washed in PB followed by double-distilled water and then contrasted in freshly prepared 1% uranyl acetate for 35 min. Sections were dehydrated through a series of graded alcohols and with propylene oxide. Afterwards, they were flat embedded in Durcupan ACM epoxy resin (14040; Electron Microscopy Sciences, Fort Washington, PA). Resin-embedded sections were polymerized at 60°C for 2 days. Sections of 65 nm were cut from the outer surface of the tissue with an ultramicrotome UC7 (Leica Microsystems Inc., Buffalo Grove, IL) using a diamond knife (Diatome, fort Washington, PA). The sections were collected on formvar-coated single slot grids and counterstained with Reynolds lead citrate. Sections were examined and photographed using a Tecnai G2 12 transmission electron microscope (Fei Company, Hillsboro, OR) equipped with a digital micrograph OneView camera (Gatan Inc., Pleasanton, CA).

### Ultrastructural analysis

Serial thin sections of dorsal striatum of the mice were used in this study. Synaptic contacts were classified according to their morphology and immuno-label and photographed at a magnification of 6800-13000×. The morphological criteria used for identification and classification of cellular components observed in these thin sections were as previously described (Zhang et al., 2015). Type I synapses, here referred as asymmetric synapses, were defined by the presence of contiguous synaptic vesicles within the presynaptic axon terminal and a thick postsynaptic density (PSD) greater than 40 nm. Type II synapses, here referred as symmetric synapses, were defined by the presence of contiguous synaptic vesicles within the presynaptic axon terminal and a thin PSD. Serial sections were obtained to determine the type of synapse. In the serial sections, a terminal or dendrite containing greater than 5 immunogold particles were considered as immuno positive terminal or dendrite. Pictures were adjusted to match contrast and brightness by using Adobe Photoshop (Adobe Systems Incorporated, Seattle, WA). The frequency of gold particles for GFP signals near to asymmetric synapses was counted from 3 mice. This experiment was successfully repeated three times.

### Photometry Data Analysis

*In vitro* photometric data were acquired in Clampex 9 and initially analyzed using Clampfit v9 or v10 (Axon Instruments/Molecular Devices), exported to Excel for organization, and plotted and analyzed in GraphPad Prism 7. The PMT readout was offset corrected to zero (with the shutter closed) before basal fluorescence (F_0_) and stimulation-induced increases in fluorescence (dF) values were obtained. Most photometric data are presented as dF/F_0_ values to compensate for differences in basal fluorescence or dLight expression differences between slices. Voltammetric data were acquired in DEMON Voltammetry Software suite (Yorgason et al., 2011), exported to Excel for organization, and plotted and analyzed in GraphPad Prism 9.

*In vivo* fiber photometry data were collected using the SPCM64 9.8 Software (Becker & Hickl). The raw photon counts were exported in ascii format for analysis with custom Python or MATLAB scripts. Briefly, the fluorescence (F_raw_) values were plotted as a function of time. Then, these fluorescence time series were converted to dF/F_0_ in two ways. For the segmented photometry data (60-s per individual trial) obtained from operant conditioning sessions, F_0_ was set to a moving average of each point before the onset of CS+, using a sliding window of +/-15s, similar to previously published methods (Patriarchi et al., 2018). Therefore, the baseline was normalized across all measurements such that only phasic dopamine transients were quantified. Occasionally, there was a negligible bleaching effect (<0.1% of F_0_ degradation) within the 60-s of acquisition in our measurement setup even without the moving average normalization. For the continuous photometry measurements over 60min duration in the *in vivo* pharmacology experiments, actual recording was performed after a 20min initial bleaching period without recording. The F_0_ was estimated by a curve fitting method. We first validated that the degradation of fluorescence intensity was well predicted by an exponential decaying model. For this validation, we used control mice virally transduced to express GFP in dorsal striatum and dLight expressing mice having no drug injection to confirm the baseline prediction method. Then, for the actual experimental animals with cocaine and saline injections, the baseline curves of F_0_ were estimated from the 20-min baseline measurement prior to the i.p. injection. Once the F_0_ baseline was estimated, dF/F_0_ was calculated using the standard method, in which dF = F_raw_-F_0_ and F_0_ is the predicted baseline by exponential curve fitting.

### Dopamine Transient Decay Analysis

For each CS+ trial, maximum peak location was identified using custom Matlab code. Then the fluorescence profile (dF/F_0_) was normalized to the maximum intensity. The data points following the maximum intensity peak were fit using a double exponential decay model. In the fitted curve, the time point where the normalized fluorescence profile passed under 36.8% of the maximum intensity was selected as the transient decay time (or lifetime) of the phasic dopamine activity in each brain regions.

### Drugs and Reagents

Cocaine-HCl was obtained from the National Institute on Drug Abuse. Dihydro-β-erythroidine hydrobromide (DHBE), nomifensine, quinpirole-HCl, and SCH23390 were purchased from Tocris Bioscience. Dopamine-HCl was purchased from Sigma Aldrich. All drugs were dissolved in ACSF. All other drugs and reagents, unless otherwise indicated, were obtained from Sigma Aldrich.

### Statistics

GraphPad Prism 9.2 was used for all data analysis and statistics. For I-O curve comparison, two way ANOVA (method and stimulation intensity) was used. For the analysis of the caspase lesion curves, a mixed effects model was used (genotype and stimulation intensity factors). For the quinpirole inhibition and high calcium experiments, unpaired two-tailed t-tests were used.

For data comparing *in vivo* transient amplitude, frequency, and decay constant changes, unpaired two-tailed t-tests were used. For analysis of the dorsal striatum subregion changes in decay constant, one way ANOVA was used. Unless otherwise indicated, all data represent the mean ± SEM.

## Acknowledgements

We would like to thank Dr. Huaibin Cai for the Casp3 virus, Dr. Joseph Cheer for his comments on the data and manuscript, Dr. Gabriel Loewinger for advice on data analysis/statistics, and Jacob Nadel and Lucas Voyvodic for their expert technical assistance with stereotaxic virus infusions. We also thank Rong Ye and Kevin Yu of the Confocal and Electron Microscopy Core, NIDA IRP, for collecting confocal and immune-electron microscopic images used in this study.

This work was supported by research grants from the National Institute on Alcohol Abuse and Alcoholism (K99R00 AA025991 to AGS, K99R00 AA027740 to SMA, and ZIA AA000407 & ZIA000416 to DML), the Intramural Research Program of the National Institute on Drug Abuse, the BRAIN Initiative (U01NS090604 and U01NS013522 to LT), and the National Institutes of Health (DP2MH107056 to LT).

## Author Contributions

Designed Research: AGS, JOL, MM, YM, DML Contributed Essential Reagents: TP, LT Performed Research: AGS, YM, JOL, SMA, SZ Analyzed Data: AGS, YM, JOL, SMA, SZ, MM Wrote the Paper: AGS, JOL, YM, DML

