## Supplementary Figures and Table for "Distinct sub-second dopamine signaling in dorsolateral striatum measured by a genetically-encoded fluorescent sensor"

### Supplementary Materials

- **Supplementary Figure1**
- **Supplementary Figure2**
- **Supplementary Figure3**
- **Supplementary Figure4**
- **Supplementary Figure5**
- **Supplementary Table1**

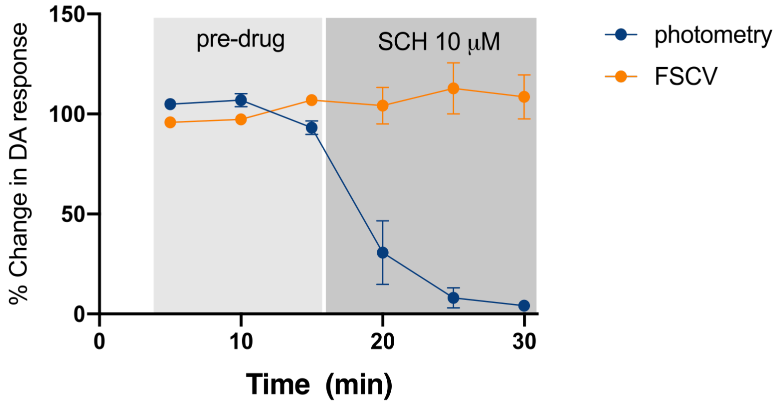


**Supplementary Figure 1**. Effect of a D1 antagonist on simultaneous FSCV and photometry measurements in brain slices expressing dLight. Comparison of responses before and after D1 antagonist application indicates that the FSCV measurement was not altered by D1 receptor blockade.

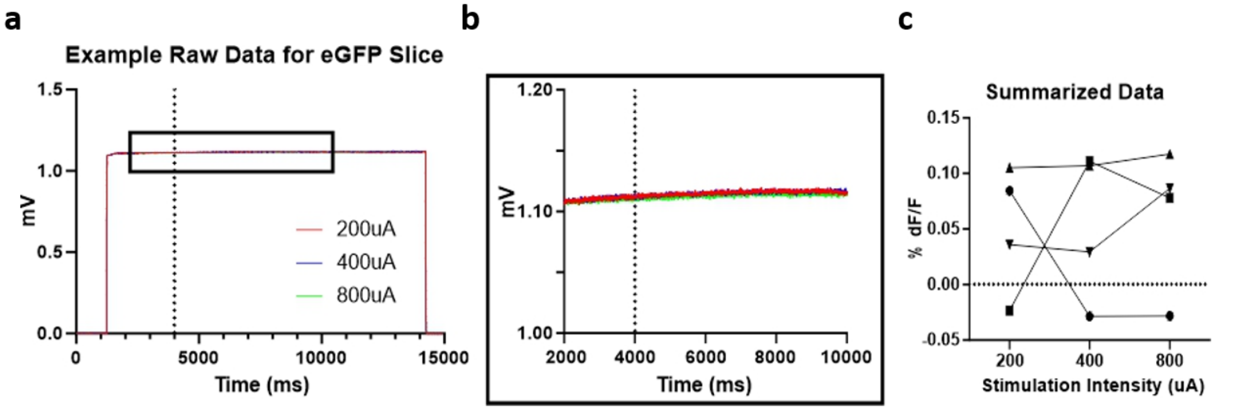


**Supplementary Figure 2**. **eGFP fluorescence does not change with electrical stimulation in brain slice experiments.** **(a)** Example of the raw data collected from a single photometry experiment. The rapid increase and decrease in signal intensity (at ~1000 msec & 1400msec) reflects the opening and closing of the fluorescent lamp shutter. The vertical dotted line indicates the time of the electrical stimulation. **(b)** Traces from panel a with an expanded y axis showing only the time when the shutter is open. **(c)** Summary data of dF/ $F_0$ at different stimulation intensity (n=4 slices). Note the lack of any consistent response to stimulation and that the scale of the fluorescence signal in the summarized data (<0.15% dF/ $F_0$) is well below any of the evoked dLight transients (typically >15% dF/ $F_0$ for slice experiments).

| | Total Number of Gold Particles | Gold Particles near asymmetric synapses | Percentage of gold particles near asymmetric synapses over the total number of gold particles (mean±SEM) | Avg. of all cases |
|---|---|---|---|---|
| Case 1 | 11,316 | 230 | 2.06 ± 0.21 | 2.43 ± 0.41 |
| Case 2 | 2,740 | 56 | 1.98 ± 0.31 | |
| Case 3 | 8,131 | 250 | 3.25 ± 0.23 | |

**Table S1. Frequency of gold particles (GFP) near to asymmetric synapses**

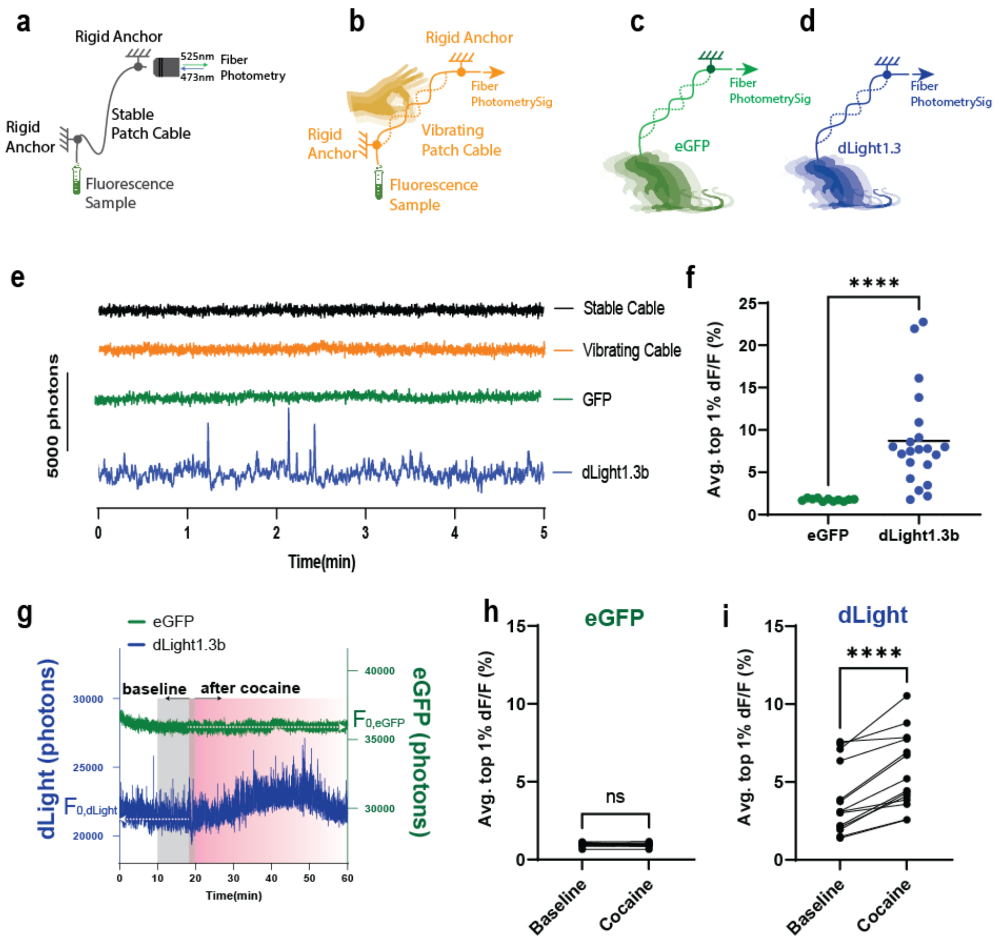


**Supplementary Figure 3**. **Cocaine-induced dLight signal is not generated from movement artifacts. (a)** Schematic diagram of the fiber system and the benchtop measurement system using a fluorescent polymer (World Precision Instruments, cat # KWIK-CAST) with a stable patch cable on an optical bench (excitation; 473nm, emission; 525nm). **(b)** Diagram of measurement with a vibrating patch cable. **(c)** Diagram of *in vivo* fiber photometry measurement with an eGFP expressing mouse. **(d)** Diagram of *in vivo* Fiber photometry measurement with a dLight expressing mouse. **(e)** Example raw traces of photon counts for 5 min from stable patch cable in the benchtop test system (black) and vibrating patch cable (orange), and for an eGFP-expressing mouse (green), and a dLight-expressing mouse (blue). Mice were freely moving in an open arena during the measurement. **(f)** Average dF/F amplitude of the photometry signals that reached the highest 1% observed during the 5 min test period for eGFP (n=11) and dLight (n=21) measured in an open field condition for 5 min ($p<0.0001$, two-tailed Mann-Whitney test). **(g)** Example raw traces of photon counts for eGFP (green, right y-axis) and dLight (blue, left y-axis) expressing mice before and after cocaine injection (15mg/kg). After 20min of initial bleaching period (without recording), the photometry recording was performed for 60min. **(h)** Average dF/F amplitude of the photometry signals that reached the highest 1% observed from eGFP expressing mice (n=11) during baseline measurement (10min average before injection) and after cocaine injection (20-30min average after injection). Two-tailed paired t-test, $p>0.05$ **(i)** Average dF/F amplitude of the photometry signals that reached the highest 1% observed from dLight expressing mice (n=14) before and after cocaine injection (two-tailed paired t-test, $p<0.0001$).

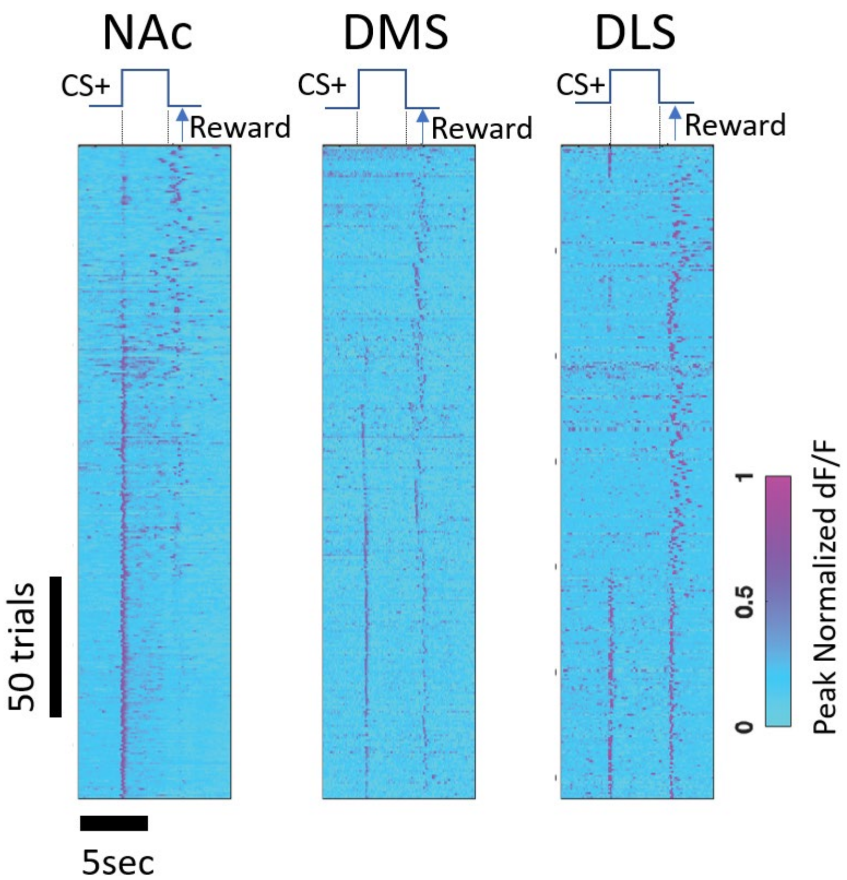


**Supplementary Figure 4**. Example DA activity patterns in NAc, DMS, and DLS across CS+ trials. The dF/F color code is normalized to the maximum peak of each trial.

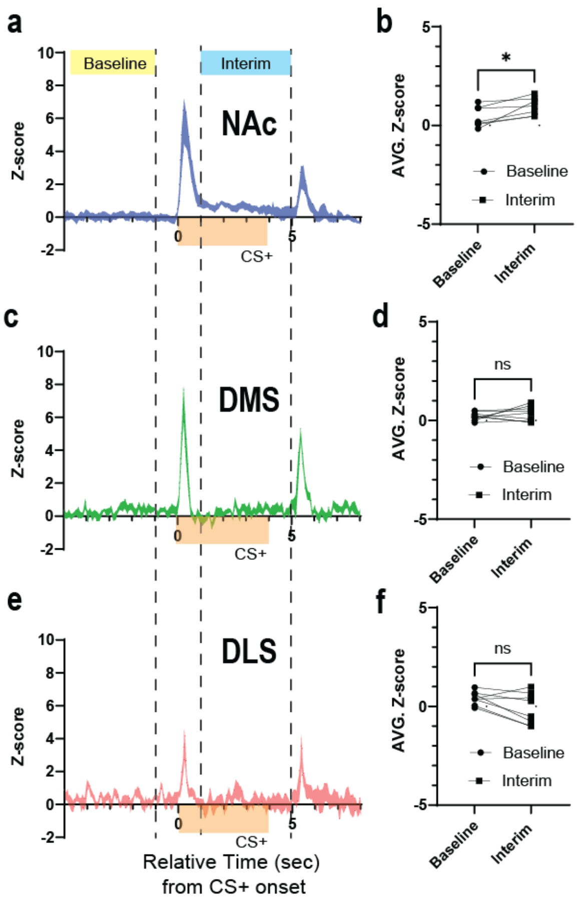


**Supplementary Figure 5.** Sustained DA increase in NAc during performance of the Pavlovian conditioning task. **(a)** Averaged dLight photometry traces of 15 CS+ trials recorded in NAc (n=6 mice) on day 14. Dashed lines indicate the end of the baseline measurement period (lefthand dashed line, -5s~-1s relative to the CS+), and the beginning and end of the "interim" period beginning after the offset of the response to the CS+ (middle dashed line, 1s after onset of CS+) and ending just before reward delivery (righthand dashed line, 5s after onset of CS+). **(b)** NAc average z-score comparison between the baseline time and interim period (two-tailed paired t-test, p=0.00361, t=2.843, df=5). **(c)** averaged dLight photometry traces of CS+ trials recorded in DMS (n=7 mice). **(d)** DMS average z-score comparison between the baseline and interim period (two-tailed paired t-test, p=0.2779, t=1.193, df=6). **(e)** averaged dLight photometry traces of CS+ trials recorded in DLS (n=7 mice) on day 14. **(f)** DLS average z-score comparison between the baseline and interim periods (two-tailed paired t-test, p=0.1134, t=1.853, df=6), Shaded areas on traces indicate SEM. Orange bar indicates CS+ duration. Note the significant increase in fluorescence in the interim period between the end of the CS+ response and reward delivery was only observed in NAc.
